# Direction-dependent neural control of finger dexterity in humans

**DOI:** 10.1101/2023.04.25.538234

**Authors:** Ohad Rajchert, Shay Ofir-Geva, Yoel Melul, Mona Khoury-Mireb, Orit Wonderman Bar-Sela, Osnat Granot, Tom Caspi, Silvi Frenkel Toledo, Nachum Soroker, Firas Mawase

## Abstract

Humans, more than all other species, skillfully flex and extend their fingers to perform delicate motor tasks. This unique dexterous ability is a product of the complex anatomical properties of the human hand and the neural mechanisms that control it. Yet, the neural basis that underlies human dexterous hand movement remains unclear. Here we characterized *individuation* (fine control) and *strength* (gross control) during flexion and extension finger movements, isolated the peripheral passive mechanical coupling component from the central neuromuscular activity involved in dexterity and then applied voxel-based lesion mapping in first-event sub-acute stroke patients to investigate the causal link between the neural substrates and the behavioral aspects of finger dexterity. We found substantial differences in dexterous behavior, favoring finger flexion over extension. These differences were not caused by peripheral factors but were rather driven by central origins. Lesion-symptom mapping identified a critical brain region for finger individuation within the primary sensory-motor cortex (M1, S1), the premotor cortex (PMC), and the corticospinal (CST) fibers that descend from them. Although there was a great deal of overlap between individuated flexion and extension, we were able to identify distinct areas within this region that were associated exclusively with finger flexion. This flexion-biased differential premotor and motor cortical organization was associated with the finger individuation component, but not with finger strength. Conversely, lesion mapping revealed slight extension-biases in finger strength within descending tracts of M1. From these results we propose a model that summarizes the distinctions between individuation and strength and between finger movement in flexion and extension, revealed in human manual dexterity.

## Introduction

Humans, extensively more than any other species, skillfully use their fingers to perform various complex tasks, from buttoning off a shirt, or tying shoelaces, up to playing musical instruments such as a saxophone. This unique ability is a product of the complex anatomical properties of the human hand and the neural mechanisms that control it, together allowing precise and versatile dexterous movement. The more sophisticated the task, the more it involves individuated finger movements, in which one or more fingers move with minimal motion of other digits or the wrist. Hand versatility primarily involves movement of the digits in two major directions, *flexion* and *extension*. In various daily activities, for example, when holding and later releasing a heavy object, task performance involves co-activation of the different fingers, first in flexion and later in extension. However, dexterous performance of certain tasks often requires execution of flexion and extension movements in individual fingers in isolation. For example, when playing the piano, isolated flexion of one finger is often needed to emit the desired musical note, and subsequent extension of that finger determines the time at which the note ends. How does the brain ensure the precision of such finger movements?

In 1968, Lawrence and Kuypers^1^ showed that bilateral pyramidal tract sectioning abolishes monkeys’ capacity for precise control of finger movement, leaving more proximal limb movements relatively less affected. Later research on finger individuation in non-human primates (NHP), as well as lesion studies in human stroke patients, implied the motor cortex and its corticospinal projections in dexterous finger movement ability^2, 3^. Despite these advances, various details concerning the neural control underlying dexterous finger flexion and extension in humans remain largely unknown. Thus, we do not know yet if the physiological mechanisms guiding finger flexion and extension depend on similar or distinct neural substrates. Evidence from functional-imaging studies suggests that finger extension and flexion movements have a shared representation in the primary motor cortex. For example, a recent carefully controlled high-resolution fMRI experiment in humans demonstrated that activation patterns in the sensorimotor hand area are highly similar when a person executes a flexion movement or an extension movement in the same finger^4^. In rhesus monkeys, neuronal activity during individuated flexion or extension of each of the five digits or the wrist is distributed throughout the same territory in M1, with little if any spatial segregation of the activity center of mass for the different movements^5^. Additionally, previous studies in macaque monkeys, have shown that the same M1 neuron may exhibit a time-locked activity prior to individuated finger flexion or extension movements, as well as prior to movements of non-adjacent digits (e.g., flexion of the thumb or middle finger, but not the index finger)^6, 7^.

Despite the above findings, there are reasons to doubt the idea of shared neural control of finger flexion and extension. First, the control of fingers’ flexion movements seems to be more precise compared to extension movements. Both in humans and monkeys, production of extension forces in fingers was found to be accompanied with higher enslaving, or unintended force production in other fingers, compared to production of flexion forces in fingers^8, 9^. It is difficult to reconcile this observation with the idea that similar neural control drives both finger extension and flexion. Yet, available data did not rule out possible effects of confounding variables, such as differences in the biomechanical properties of movements in the two directions, or soft tissue coupling between finger extension and flexion, nor investigated finger individuation while controlling for differences in strength across movement directions. Second, generalization between flexion and extension following multiday training is asymmetrical. That is, learning extension movements generalized to flexion, but not vice versa^10^. Third, micro-stimulation over the human motor cortex evoked flexion of the fingers far more often than extension^11^. While this last observation may suggest that flexion is more extensively represented in the motor cortex, it may also be explained by the fact that the representation of finger extensor movements in M1 is more buried in the anterior bank of the central sulcus, thus is relatively less accessible to electrical stimulation. Lastly, people with hemiparesis due to stroke affecting M1 or the corticospinal tract, most often exhibit more severe weakness in extension compared to flexion movements of the fingers^12^.

Past human lesion studies aiming to unravel the neural basis of dexterous distal upper-limb movement often failed to fully dissociate force generation and individuation capacities. Moreover, testing was often done late after stroke onset, thus obscuring the natural physiological picture by effects of neural plasticity and adoption of compensatory movement strategies. The aim of the current study was to determine whether the neurophysiological mechanisms that underlie motor control of finger flexion are distinct from those that govern finger extension. We began by questioning whether the characteristics of dexterous finger flexion are different from those of finger extension, in healthy people. Next, we asked whether mechanical coupling between digits contributed to movement of uninstructed fingers and thereby limited the ability to individuate finger movement. Lastly, we applied a voxel-based lesion-symptom mapping, to explore the neural substrates critical for individuation capacity and force generation in fingers’ flexion vs. extension, in a group of early sub-acute stroke patients with upper-limb paresis.

## Results

### Behavioral evidence for distinct control of finger flexion and extension

In Experiment 1, we aimed to characterize the behavioral principles underlying the control process of single and multi-digit dexterous motor activity in the flexion and extension directions, through a novel finger dexterity task. Healthy young participants (n=13, age: 28.2±2.9 years) performed three isometric tasks: finger strength, single-finger individuation and multi-finger individuation. In the *finger strength task*, we measured for each digit the maximal forces in the flexion and extension directions (MVF; maximal voluntary contraction force). We defined the ***Strength*** of each digit by calculating the 95th percentile of the force traces produced across all sampled force data points during the finger-pressing period in each trial and then averaged across the two MVF trials (See Methods). The overall strength (i.e., hand level strength) was then calculated by averaging the Strength index across all 5 digits. Of note, due to limitations in the force capability of the robot’s sensors, participants were allowed to produce forces only within the dynamic range (±20 N, positive values for extension and negative for flexion). When the produced force exceeded these values, a warning signal appeared, and participants were instructed not to pass this limit. Out-of-range forces were automatically floored to ±20 N (see Discussion). As predicted, participants applied significantly more force during finger flexion compared to extension as revealed by a two-way RM-ANOVA, which showed significant effects for the direction (𝐹_1,12_ = 882.05, 𝑝 = 1.33 × 10^−1^^2^) and the digit (𝐹_4,48_ = 14.10, 𝑝 = 1.11 × 10^−7^), though non-significant interaction between the two factors (𝐹_4,48_ = 2.16, 𝑝 = 0.08) (Fig. 1H).

**Figure 1.**
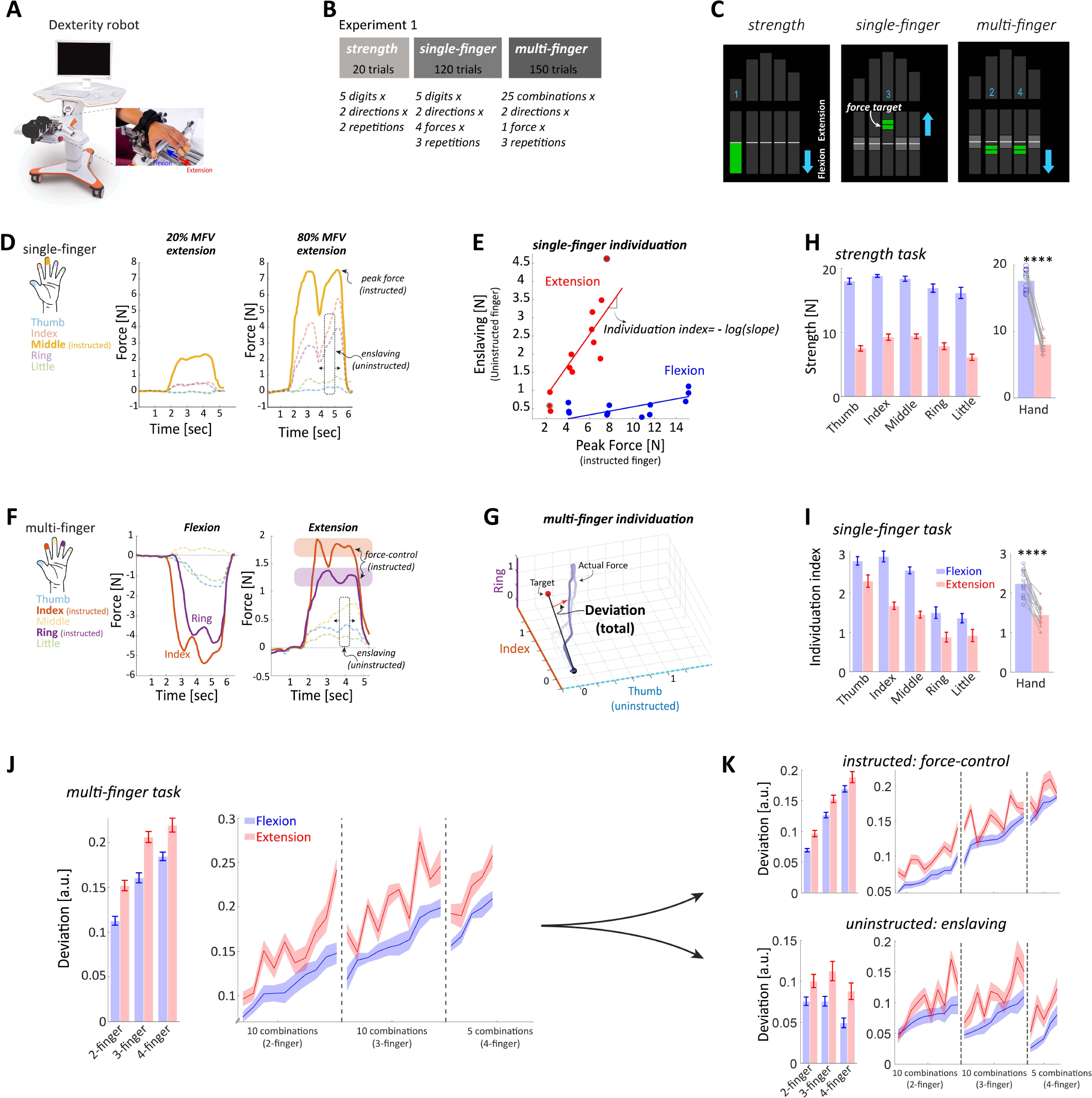
– Experiment 1 setup, experimental protocol and results. (A) The dexterity robot (Amadeo, Tyromotion®) used during all conducted experiments. On the right, the physical connection between the robot’s handles and the tested hand’s fingers via magnets. (B) The protocol of experiment 1, partitioned into three tasks: strength, single finger individuation and multi-finger individuation. (C) The Graphic User Interface (GUI) used during all experiments. The diagram on the left shows the GUI during a strength task, specifically describing the case in which participants were asked to produce their maximal force using the thumb in the flexion direction. The middle diagram shows the GUI during a single-finger individuation task, specifically describing the case in which participants were asked to extend their middle finger to 60% of the corresponding maximal force. The diagram on the right shows the GUI during a multi-finger individuation task, specifically describing the case in which participants were asked to flex both their index and ring fingers simultaneously to 25% of the corresponding maximal forces. (D) Examples of extension trials from the single-finger individuation task, specifically middle finger movements. Target forces were set to be 20% and 80% of the corresponding maximal force in the left and right traces, respectively. The peak force property of the instructed finger and the enslaving property of the uninstructed fingers are indicated by arrows. (E) Linear regression graph contains 12 extension (red) and 12 flexion (blue) trials (3 repetitions of 4 force levels) of a single finger. The two examples brought before are highlighted in gray circles. Individuation index of each finger in a certain direction is calculated as the -log of the slope between the peak force of the instructed finger and the enslaving of the uninstructed fingers. (F) Examples flexion and extension trials from the multi-finger individuation task, specifically index and ring fingers movements. Flexion forces were assigned negative values and extension forces had positive value. The force control component of the instructed fingers and the enslaving component of the uninstructed fingers are indicated by arrows. (G) A 3D illustration of the deviation which occurs during multi-finger movements. The total deviation between the ideal “init-to-target” projection and the actual force trace is taking into consideration all fingers together. (H). Strength values (mean ± SE) obtained in the strength task. Finger-level group data and hand-level individual data are presented on the left and right graphs, respectively. (I) Individuation index values (mean ± SE) obtained in the single-finger individuation task. Finger-level group data and hand-level individual data are presented on the left and right graphs, respectively. (J) Total deviation values (mean ± SE) obtained in the multi-finger individuation task. Chord-type group data and all combinations sorted in ascending order are presented on the left and right graphs, respectively. (K) top - Instructed deviation values (mean ± SE) obtained in the multi-finger individuation task. Chord-type group data and all combinations sorted in ascending order are presented on the left and right graphs, respectively. bottom - Uninstructed deviation values (mean ± SE) obtained in the multi-finger individuation task. Chord-type group data and all combinations sorted in ascending order are presented on the left and right graphs, respectively. Bars represent mean and Errors bars represent ±SEM. Dots are individuals. **** p<0.0001.

In the *single-finger individuation task*, participants were instructed to exert force, each time with a different single digit, either in flexion or extension, to maintain one of four target force levels (20, 40, 60 and 80% of the MVF), while keeping all other digits as static as possible. Each digit’s ability to produce independent movements was quantified by the ***individuation index*** (II), defined as the degree to which all uninstructed digits remain still during movement of a particular instructed digit. The higher this index is, the better the independence capability (Fig. 1D-E). We found that digit flexion generally exhibited superior strength and independence compared to digit extension, providing strong evidence for the direction-dependent control hypothesis of single-digit dexterous actions. Our data showed that the individuation index in the extension direction was significantly lower than the flexion direction (Fig. 1I). Two-way RM-ANOVA with factors of direction and digit revealed significant effects for the direction (𝐹_1,12_ = 79.32, 𝑝 = 1.23 × 10^−6^), the digit (𝐹_4,48_ = 64.82, 𝑝 = 9.47 × 10^−1^^9^), as well as significant direction × digit interaction (𝐹_4,48_ = 7.72, 𝑝 = 6.90 × 10^−5^) (Fig. 1F and Supplementary material).

During the *multi-finger individuation task*, participants (n=12), performed 25-digit combinations (“chords”; divided into 3 types: 10 two-digit, 10 three-digit and 5 four-digit), to maintain one force level (25% of MVF) with the instructed digits in each direction, while keeping the uninstructed digits as immobile as possible. The ability to generate accurate multi-digit force combinations was separated into two aspects – force control, calculated as the deviation of the instructed digits from the target (***DeviationI***), and enslaving, calculated as the deviation of the uninstructed digits from zero (***DeviationU***). These deviations were combined into a complete measurement taking into consideration all digits (***DeviationT***). The lower these deviations are, the better the control level (Fig. 1F-G).

Our results show that total accuracy (*DeviationT*) during the multi-finger task was reduced for finger extension compared to flexion. Two-way RM-ANOVA with factors direction and chord type showed significant effects for the direction (𝐹_1,11_ = 58.49, 𝑝 = 9.99 × 10^−6^), chord type (𝐹_2,22_ = 129.49, 𝑝 = 6.78 × 10^−1^^3^), but no significant direction × chord type interaction (𝐹_2,22_ = 1.03, 𝑝 = 0.37) (Fig. 1J). When considering the instructed fingers alone (*DeviationI*), we found that instructed fingers reach more accurately the target force in the flexion direction compared to extension direction, as revealed by Two-way RM-ANOVA with significant effects of direction (𝐹_1,11_ = 23.14, 𝑝 = 5.44 × 10^−4^) and chord type (𝐹_2,22_ = 446.58, 𝑝 = 1.55 × 10^−1^^8^), with non-significant interaction between the two (𝐹_2,22_ = 0.95, 𝑝 = 0.40) (Fig. 1K). Improved accuracy in finger flexion over extension movement was also evident when investigating the enslaving, i.e., the deviation of the uninstructed fingers from the baseline force level (*DeviationU*). Two-way RM-ANOVA showed significant effects for the direction (𝐹_1,11_ = 18.12, 𝑝 = 0.001) and chord type (𝐹_2,22_ = 22.10, 𝑝 = 5.47 × 10^−6^), though only trend for interaction between the two (𝐹_2,22_ = 3.01, 𝑝 = 0.07) (Fig. 1K).

In summary, our behavioral assessments in healthy subjects show that finger movements have more strength and larger individuation capability in the flexion direction compared to extension. In addition, participants were more accurate in both controlling forces of the instructed finger(s) and immobilizing the uninstructed finger(s) in the flexion direction, regardless of the exerted force.

### Contribution of mechanical coupling to behavioral differences shown in finger flexion and extension

The above behavioral measures of finger dexterity may reflect not only the impact of neural control mechanisms but also of mechanical properties of the effector organ (Fig. 2A). In Experiment 2 we sought to assess the extent to which coupling between fingers, reflecting mechanical properties the hand’s soft tissues and tendons, contribute to the movements of uninstructed digits (i.e., enslaving). Healthy young participants (n=20, age 31.7±6.1 years) underwent a task in which each digit was moved *passively* by the robot in flexion and extension directions, while measuring the isometric forces created concurrently in the unmoved digits (Fig. 2B). Participants were asked to remain at rest during 2 blocks in which the robot moved each digit: (1) through four distances (20, 40, 60 and 80% of the Range of Motion; *ROM* block) at the maximal possible velocity (0.2 m/s) and (2) at three velocities (0.05, 0.1, 0.15 m/s, *Velocity* block) through a single distance (set to 80% of the ROM) (Fig. 2C). The involuntary forces created in the unmoved digits (i.e., enslaving) were recorded during the task and used for quantifying the mechanical coupling component (Fig. 2D-E). The lower the enslaving in this task, the less the impact of hand anatomy on finger dexterity measures.

**Figure 2.**
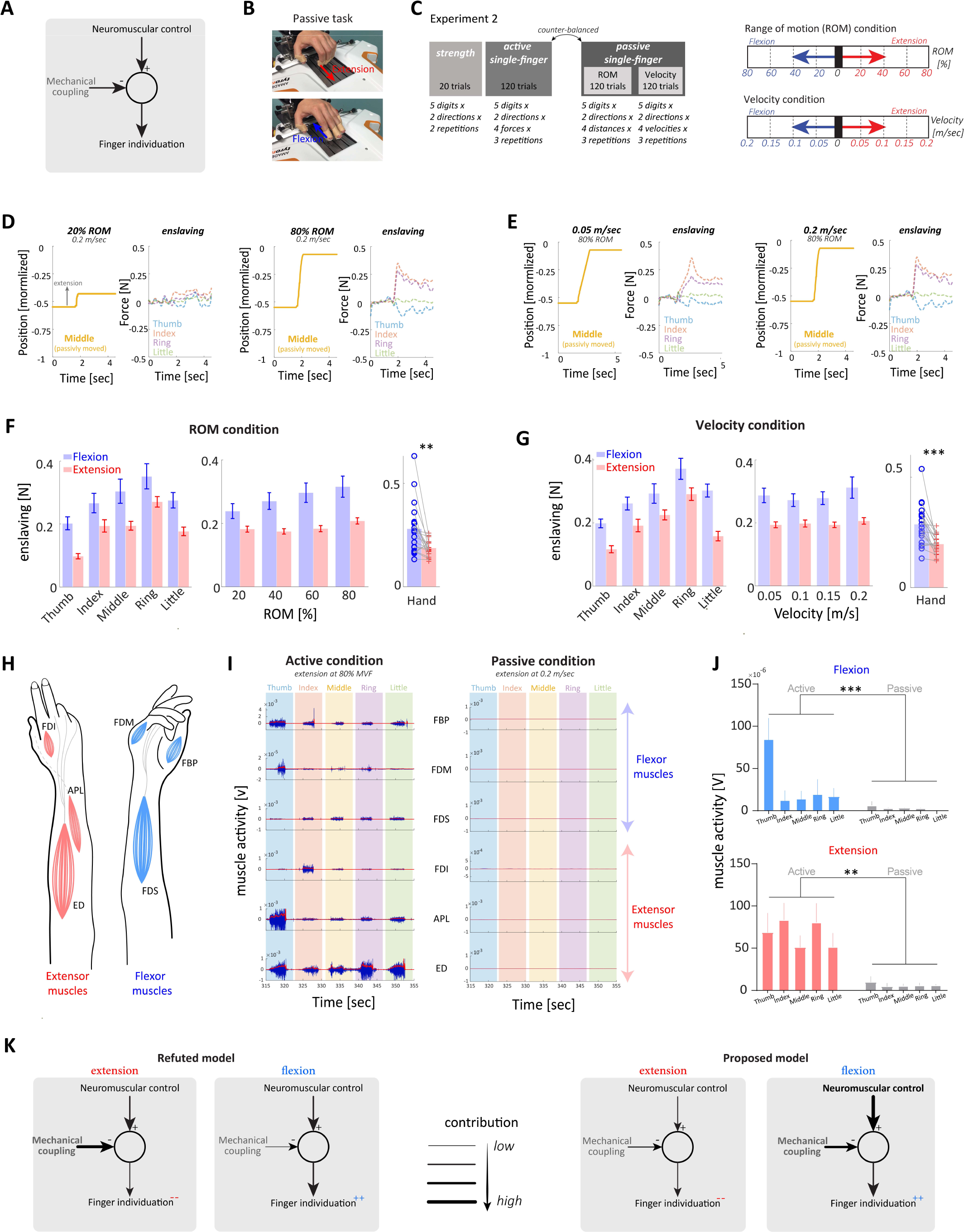
– Setup, experimental protocol and findings of Experiment 2. (A) General model regarding the two main contributors to the finger individuation ability that were focused on in this study: neuromuscular control and mechanical coupling. (B) Setup. The two different working positions in the passive task depends on the direction of the passive movement (flexion/extension). (C) Protocol. On the left, the protocol of experiment 2, partitioned into two counter-balanced parts: active contains the strength and single-finger individuation tasks and passive contains two conditions: Range of Motion (ROM) and velocity. On the right, ROM condition options (20, 40, 60 and 80% of ROM) and velocity condition options (0.05, 0.1, 0.15, 0.2 m/s). (D) Examples of middle finger extension trials from the passive task during the ROM condition. Specifically, movement’s velocity was set to be constant at 0.2m/s, while distance varied between 20% and 80% of the ROM in the left and right traces, respectively. The enslaving property of the uninstructed fingers forces was recorded during the passive movement. (E) Examples of middle finger extension trials from the passive task during the velocity condition. Specifically, movement’s distance was set to be constant at 80% of the ROM, while velocity varied between 0.05 and 0.2 m/s in the left and right traces, respectively. The enslaving property of the uninstructed fingers forces was recorded during the passive movement. (F) Enslaving values (mean ± SE) obtained in the passive task during the ROM condition. Finger-level group data is presented on the left graph, ROM-level group data is presented on the middle graph and hand-level individual data is presented on the right graph. (G) Enslaving values (mean ± SE) obtained in the passive task during the velocity condition. Finger-level group data is presented on the left graph, velocity-level group data is presented on the middle graph and hand-level individual data is presented on the right graph. (H) Schematic illustration of recorded digit flexor and extensor muscles. FPB-flexor pollicis brevis, FDM-Flexor digiti minimi brevis, FDS-flexor digitorum superficialis, FDI-first dorsal interosseous, APL-abductor pollicis longus, ED-extensor digitorum. (I) Example of muscle activity in one trial in one participant during active (extension at 80% MVF trial) and passive condition (extension at vel of 0.2m/sec). The upper 3 rows show data for the digit flexor muscles and the lower 3 rows show the data of the digit extensor muscle. (J) Average muscle activity envelope across participants in each direction and digit across active and passive conditions. (k) Our data refuted the model (left) suggesting similar neuromuscular control between directions and different mechanical coupling bias toward the flexion direction, together resulting in larger finger individuation capability of flexion compared to extension. Instead, we propose a new model (right) that suggests different neuromuscular control favoring of flexion rather than extension, compensating for the different mechanical coupling bias toward the flexion direction, together resulting in larger finger individuation capability in flexion compared to extension. Arrows’ thickness reflects the strength of the contribution of each component where thin lines represent weaker and thick lines represent stronger involvement. Bars represent mean and Errors bars represent ±SEM. Dots are individuals. ** p<0.01 and *** p<0.001.

We found that enslaving due to mechanical coupling is much larger when the robot passively flexes the finger than when it extends the fingers (Fig. 2F). Two-way RM-ANOVA with direction and digit factors revealed significant effects for the direction (𝐹_1,19_ = 12.62, 𝑝 = 0.002) and the digit (𝐹_4,76_ = 29.20, 𝑝 = 1.04 × 10^−1^^4^), but not for the interaction between the two (𝐹_4,76_ = 0.48, 𝑝 = 0.75). When considering the mechanical coupling across the different ROM ranges, we found significant effects for the direction factor (𝐹_1,19_ = 12.62, 𝑝 = 0.0021) and the range factor (𝐹_3,57_ = 15.43, 𝑝 = 1.83 × 10^−7^), as well as an interaction between the two (𝐹_3,57_ = 10.77, 𝑝 = 1.05 × 10^−5^). Overall, this result indicated that mechanical coupling was significantly lower for extension compared to flexion direction. The *Velocity* condition also showed that the contribution of mechanical coupling to enslaving is much larger in the flexion direction (Fig. 2G). Two-way (direction vs digit) RM-ANOVA revealed significant effects for the direction (𝐹_1,19_ = 22.79, 𝑝 = 1.32 × 10^−4^) and the digit (𝐹_4,76_ = 31.58, 𝑝 = 1.72 × 10^−1^^5^), with marginal trend for interaction between the two (𝐹_4,76_ = 2.22, 𝑝 = 0.07). When examining whether enslaving is affected differently in the different robot velocities, we found an effect only for the direction factor (𝐹_1,17_ = 20.81, 𝑝 = 2.77 × 10^−4^), with no effect for the velocity factor (𝐹_3,51_ = 2.08, 𝑝 = 0.11) and no interaction between the two (𝐹_3,51_ = 0.70, 𝑝 = 0.56).

We confirmed using EMG that enslaving forces measured in Experiment 2 were unquestionably the consequence of mechanical coupling between the digits, and not a result of muscle contraction in response to passive stretching by the robot. In an additional cohort of participants (n=7, age 36.3±7.9 years), we recorded EMG from 6 hand muscles - flexor pollicis brevis (FPB), flexor digiti minimi brevis (FDM), flexor digitorum superficialis (FDS), first dorsal interosseous (FDI), abductor pollicis longus (APL), extensor digitorum (ED) - when participants actively made finger flexion and extension and when the robot passively moved the fingers (Fig. 2H-I). EMG activity during the passive condition was significantly lower than the activity during the active conditions (Fig. 2I-J). Mixed-effect model in the flexion direction with condition (active, passive) and digit factors revealed significant effects for the condition (𝐹_1,6_ = 38.45, 𝑝 = 0.0008) and the digit (𝐹_4,24_ = 33.30, 𝑝 < 0.0001), and significant condition × digit interaction (𝐹_4,17_ = 26.75, 𝑝 < 0.0001). Mixed-effect model in the extension direction also revealed a significant effect of the condition (𝐹_1,6_ = 26.91, 𝑝 = 0.002), but no significant digit effect (𝐹_4,24_ = 0.56, 𝑝 = 0.69), nor condition × digit interaction (𝐹_4,17_ = 0.58, 𝑝 = 0.68).

Altogether, these results suggest that digit flexion is more affected than digit extension by soft-tissue biomechanical factors. Apparently, these factors would have limited individuation of finger flexion to a larger extent compared to finger extension. Nevertheless, healthy subjects were found to possess much larger individuation capacity in the flexion direction. Thus, our findings rule out the possibility that increased mechanical coupling accounted for the reduced individuation capacity in extension compared to the flexion. Rather, they suggest that the neural mechanisms enabling dexterous finger movements in humans reached a higher level of sophistication in control of finger movements performed in the flexion direction compared to the extension direction.

### Stroke lesion impact on finger dexterity

In experiment 3, our goal was to investigate the causal effect of cortical and sub-cortical lesions on finger dexterity impairment. We quantified individuation capacity and strength in finger flexion and extension movements, in a cohort of first-event stroke patients (n=32, age: 60.5±10.6 years, time after stroke onset: 40.9±17.0 days; see Table 1) who performed the finger strength and single-finger individuation tasks using their paretic hand (Fig. 3A-B). Patients’ data was compared to that of an age-matched healthy control group (n=11, age: 56.3±10.0 years) who performed the same tasks with both the right and left hands (the average performance between hands was used as an index value for comparison with patients’ performance). In the stroke group, individuation capacity was significantly lower for finger extension compared to flexion (Fig. 3C-D). A two-way (direction, digit) RM-ANOVA in this group revealed a significant effect for the direction factor (𝐹_1,31_ = 30.22, 𝑝 = 5.16 × 10^−6^) reflecting flexion advantage, and the digit factor (𝐹_4,124_ = 25.43, 𝑝 = 2.15 × 10^−1^^5^), with an interaction between the two (𝐹_4,124_ = 14.37, 𝑝 = 1.15 × 10^−9^). In the control group, individuation was also significantly lower for extension compared to flexion direction, as revealed by a two-way RM-ANOVA with significant effects for direction (𝐹_1,10_ = 82.41, 𝑝 = 3.83 × 10^−6^) and digit (𝐹_4,40_ = 43.73, 𝑝 = 4.30 × 10^−1^^4^) factors, but no interaction between the two (𝐹_4,40_ = 1.95, 𝑝 = 0.12).

**Table 1.** –stroke patients characteristics (Experiment 3)

| Participant | Gender | Age<br>[years] | Lesion<br>type | Paretic<br>hand | Time<br>after<br>onset<br>[days] | ARAT | FMA-UE |
| --- | --- | --- | --- | --- | --- | --- | --- |
| 0102 | F | 52 | I | L | 34 | 39 | 57 |
| 0103 | M | 65 | I | L | 33 | 19 | 32 |
| 0104 | F | 65 | I/H | R | 30 | 24 | 40 |
| 0105 | M | 43 | H | R | 40 | 41 | 44 |
| 0106 | M | 74 | I | R | 49 | 39 | 53 |
| 0107 | M | 66 | I | L | 66 | 36 | 43 |
| 0108 | M | 57 | I | L | 32 | 32 | 40 |
| 0109 | F | 65 | I | L | 50 | 37 | 42 |
| 0110 | F | 56 | I | R | 50 | 57 | 62 |
| 0111 | F | 58 | H | R | 13 | 49 | 61 |
| 0112 | M | 67 | I | R | 45 | 55 | 52 |
| 0113 | M | 50 | I | R | 79 | 16 | 34 |
| 0114 | M | 63 | I/H | L | 45 | 52 | 50 |
| 0115 | F | 50 | I | R | 19 | 47 | 55 |
| 0116 | M | 34 | H | R | 49 | 57 | 65 |
| 0117 | M | 65 | I | L | 47 | 6 | 24 |
| 0118 | M | 69 | I | L | 30 | 34 | 49 |
| 0119 | M | 49 | I | R | 43 | 23 | 40 |
| 0120 | M | 67 | I | L | 52 | 52 | 58 |
| 0121 | M | 67 | I | R | 51 | 50 | 51 |
| 0123 | F | 70 | I | L | 27 | 52 | 58 |
| 0124 | M | 51 | I | R | 34 | 19 | 25 |
| 0125 | M | 70 | I | L | 55 | 23 | 41 |
| 7001 | M | 55 | I | R | 78 | 4 | 21 |
| 0126 | M | 74 | I | R | 60 | 57 | 58 |
| 0127 | M | 72 | I | R | 19 | 21 | 38 |
| 0129 | M | 78 | I | L | 17 | 23 | 36 |
| 0131 | F | 66 | H | L | 29 | 45 | 55 |
| 0132 | F | 42 | I | L | 32 | 57 | 61 |
| 7003 | M | 56 | I/H | R | 59 | 51 | 41 |
| 0134 | M | 70 | I | R | 20 | 42 | 48 |
| 0135 | M | 49 | I | R | 21 | 37 | 38 |
| mean ± STD |  | 60.5 ±<br>10.6 |  |  | 40.9 ±<br>17.0 | 37.4 ±<br>15.6 | 46.0 ±<br>11.7 |

Gender (M, male; F, female), age (years), lesion type (I, ischemic; H, hemorrhagic), paretic hand (L, left; R, right), Action Research Arm Test (ARAT; maximal score 57), Fugl-Meyer Assessment scale for the upper extremity (FMA-UE; maximal score 66).

**Figure 3.**
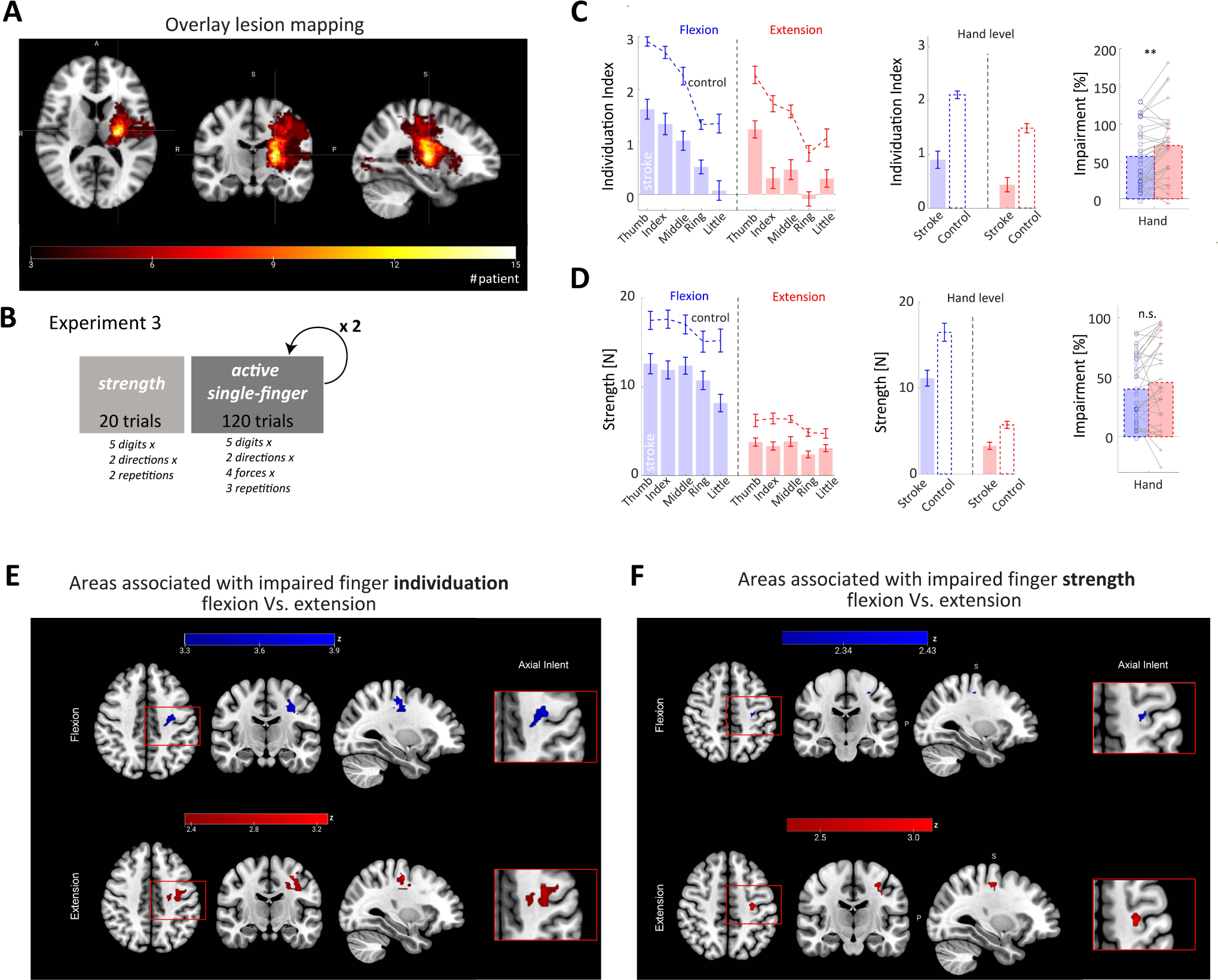
– Stroke lesion impact on finger dexterity-Experiment 3. (A) Overlay lesion mapping of stroke patients. Color bar represents the number of patients having a lesion in a voxel. (B) The protocol of experiment 3, stroke patients performed the strength and single-finger individuation tasks on the paretic hand, whereas healthy controls performed these tasks on both their right and left hand. (C) Individuation index values (mean ± SE) obtained in the single-finger individuation task. Stroke group data is depicted in full bars while control group data is depicted in doted lines (or empty bars). Finger-level groups data is presented on the left graph, hand-level groups data is presented on the middle graph and hand-level impairment individual data is presented on the right graph. (D) Strength values (mean ± SE) obtained in the strength task. Stroke group data is depicted in full bars while control group data is depicted in doted lines (or empty bars). Finger-level groups data is presented on the left graph, hand-level groups data is presented on the middle graph and hand-level impairment individual data is presented on the right graph. (E) VLSM analysis depicting voxels in which the existence of damage was associated with significant impairment of finger individuation in flexion (blue) and extension (red). For the flexion direction, voxels that survived FDR correction (p-FDR < 0.05; Z > 3.32) are shown. In the extension direction, the depicted voxels survived lenient criterion of p < 0.01 (Z > 2.32). (F). VLSM analysis depicting voxels in which the existence of damage was associated with significant reduction of finger strength in flexion (blue) and extension (blue). No voxels survived FDR correction, but a more lenient criterion of p < 0.01 (Z > 2.32). Bars represent mean and Errors bars represent ±SEM. Dots are individuals. ** p<0.01 and n.s. for not significant.

To assess finger dexterity in stroke patients relative to healthy controls, we examined individuation capacity and strength in the hand level (averaged across digits). Stroke patients had a significantly lower individuation index compared to healthy controls. Unbalanced two-way (direction, group) ANOVA revealed a significant effect for the direction factor (𝐹_1,82_ = 9.75, 𝑝 = 0.0025) reflecting advantage for flexion, and the group factor (𝐹_1,82_ = 40.86, 𝑝 = 9.41 × 10^−9^) indicating stroke-group disadvantage, with no interaction between the two factors (𝐹_1,82_ = 0.19, 𝑝 = 0.66). To assess the impact of stroke on the magnitude of finger-dexterity impairment in each movement direction, we used a normalized ***Impairment*** measure (See Methods). Finger individuation impairment was significantly larger (𝑝𝑎𝑖𝑟𝑒𝑑 𝑡 − 𝑡𝑒𝑠𝑡, 𝑡_31_ = − 3.14, 𝑝 = 0.0037) in extension (impairment of 71.17±8.17%) compared to flexion direction (56.62±7.30%) (Fig. 3C). Of note, impairment of values higher than 100% means that, relative to the controls, individuation index in stroke patients was negative (this is possible when patients applied higher forces in the uninstructed fingers than the exerted force of the instructed finger).

Finger strength was also significantly lower in extension compared to flexion, both in the stroke and the control groups. RM-ANOVA in the stroke group revealed significant effect for the direction (𝐹_1,31_ = 194.15, 𝑝 = 6.85 × 10^−1^^5^), digits (𝐹_4,124_ = 16.39, 𝑝 = 8.33 × 10^−1^^1^), and interaction between the two factors (𝐹_4,124_ = 10.13, 𝑝 = 3.98 × 10^−7^). In the control group, RM-ANOVA revealed significant effect for the direction (𝐹_1,10_ = 164.25, 𝑝 = 1.57 × 10^−7^), and the digit (𝐹_4,40_ = 9.19, 𝑝 = 2.32 × 10^−5^) but no interaction between the two factors (𝐹_4,40_ = 0.85, 𝑝 = 0.50). Unbalanced two-way (direction, group) ANOVA comparing strength at the hand level in flexion and extension between the stroke and the control groups revealed significant disadvantage for the stroke group with significant effect for the direction factor (𝐹_1,82_ = 105.49, 𝑝 = 2.18 × 10^−1^^6^), and the group (𝐹_1,82_ = 17.17, 𝑝 = 8.27 × 10^−5^), but without interaction between the two (𝐹_1,82_ = 2.45, 𝑝 = 0.12), showing that the advantage in flexion is common to stroke victims and healthy subjects alike. However, assessment of the magnitude of force-generation (fingers’ strength) *Impairment* in the stroke group did not show a significant difference between impairment in flexion and impairment in extension (𝑝𝑎𝑖𝑟𝑒𝑑 𝑡 − 𝑡𝑒𝑠𝑡, 𝑡_30_ = − 1.37, 𝑝 = 0.18); in extension (impairment of 43.52±6.81%), compared to flexion direction (38.62±4.97%) (Fig 3D).

The impact of lesion topography on finger dexterity was analyzed using voxel-based lesion-symptom mapping (VLSM)^13^. To avoid spurious results, only voxels involved in 15% or more of the patients were analyzed^14^. Given that most patients had a stroke restricted to the territory of the middle cerebral artery (MCA), such voxels were found in cortical and sub-cortical regions involved in motor control, including the primary sensory and primary motor cortices (S1, M1), premotor cortex, corona radiata, internal capsule and basal ganglia^15^, as well as adjacent, regions - inferior parietal lobule, superior temporal and inferior frontal gyri and the insula (Fig. 3A).

#### Impact of lesion topography on finger individuation

VLSM revealed two partially overlapping voxel clusters in which the existence of damage impaired individuation index for flexion and extension (Fig. 3E; Table 2). The voxel cluster associated with significantly impaired individuation in the flexion direction included 220 voxels (FDR-corrected, z-score threshold = 3.32) with peak z-score (z = 3.89) situated within the subdivision of the corticospinal tract originating in M1 (M1-CST, 21% of the cluster; MNI coordinates: -30 -22 48) and impacted also the second division of the superior longitudinal fasciculus (SLF-II, 45% of cluster), subdivision of the CST originating in the premotor cortex (PM-CST, 10% of the cluster), the superior thalamic radiation (STR, 7% of the cluster), M1 (6% of the cluster), the cortico-reticular pathway and corticospinal fibers originating in S1 (CRP and S1-CST, respectively, each comprising 3% of the cluster). No voxels associated with impaired individuation in the extension direction survived the FDR correction for multiple comparisons. We therefore report here for the extension analysis voxels that survived a lenient criterion of p < 0.01 (corresponding to minimal z-score of 2.32). For a similar approach see ^14, 16–18^. The cluster peak z-score (z = 3.26) was situated within M1-CST (8% of the cluster; MNI coordinates: -28 –16 48), while its periphery also overlapped with SLF-II (30%), M1 (19%), CRP (14%), and PM-CST (5%).

**Table 2.** VLSM analysis. Brain structures in which more than 5 significant voxels were detected by VLSM for the finger-individuation (expressed by individuation index, II) and finger-strength (expressed by normalized maximal voluntary force, MVF) at flexion and extension. *For flexion direction, voxels that survived FDR correction (p-FDR < 0.05), corresponding with Z > 3.32. are shown. For extension direction no voxels survived FDR-correction for multiple comparisons but survived a more lenient criterion of p < 0.01, corresponding with Z > 2.32. Brain structures: SLF-II = 2^nd^ division of the superior longitudinal fasciculus; M1 = primary motor cortex (without lower-limb and trunk); M1-CST, PMd-CST = corticospinal tract fibers originating in M1 and the dorsal premotor cortex, respectively; STR = superior thalamic radiation; CRP = cortico-reticular pathway.

|  |  |  |  |  |  | p < 0.01 criterion |  | P<sub>FDR</sub> < 0.05 criterion |  |
| --- | --- | --- | --- | --- | --- | --- | --- | --- | --- |
| Test | Structure | max. Z-value | X | Y | Z | Voxels | % area | voxels | % area |
| II, Flexion* | SLF-II | 3.89 | -34 | -18 | 36 | 151 | 5.52 | 98 | 3.58 |
|  | M1-CST | 3.89 | -30 | -20 | 44 | 86 | 8.49 | 46 | 4.55 |
|  | PMd-CST | 3.54 | -32 | -10 | 34 | 31 | 6.63 | 19 | 4.07 |
|  | STR | 3.32 | -28 | -16 | 36 | 63 | 1.61 | 16 | 0.41 |
|  | M1 | 3.89 | -34 | -18 | 38 | 85 | 5.59 | 14 | 1.56 |
|  | CRP | 3.32 | -24 | -22 | 38 | 68 | 2.31 | 7 | 0.24 |
|  | S1-CST | 3.32 | -28 | -20 | 36 | 19 | 2.87 | 6 | 0.91 |
| II, Extension | SLF-II | 2.92 | -38 | -12 | 30 | 105 | 3.84 |  |  |
|  | M1 | 3.26 | -32 | -18 | 50 | 66 | 7.36 |  |  |
|  | CRP | 2.45 | -24 | -18 | 34 | 50 | 1.69 |  |  |
|  | STR | 2.45 | -22 | -16 | 34 | 45 | 1.15 |  |  |
|  | M1-CST | 2.58 | -30 | -20 | 44 | 29 | 2.87 |  |  |
|  | PMd-CST | 2.92 | -32 | -10 | 34 | 12 | 2.57 |  |  |
| normalized MVF, Flexion | M1 | 2.43 | -28 | -20 | 48 | 6 | 0.59 |  |  |
| normalized MVF, Extension | SLF-II | 2.58 | -26 | -18 | 46 | 24 | 0.88 |  |  |
|  | M1-CST | 3.08 | -28 | -20 | 48 | 12 | 1.19 |  |  |
|  | M1 | 3.08 | -32 | -18 | 48 | 7 | 0.78 |  |  |

#### Impact of lesion topography on finger strength

When applying VLSM on normalized MVF data (which served as a surrogate marker of fingers’ strength), no voxels survived the FDR correction. Therefore, we report here voxels that surpassed a lenient criterion of p < 0.01 (z-score > 2.32). In the flexion direction such voxels comprised a small single cluster of 10 voxels (zmax = 2.43, MNI coordinates –28 –20 48), of which 6 voxels (60%) were located in M1-CST. In the extension direction a larger cluster of 66 voxels (zmax = 3.08, MNI coordinates –28 –20 48) emerged. This cluster involved the SLF-II (50%), M1-CST (18%) and M1 (11%) (Fig. 3F; Table 2). The cluster associated with flexion strength overlapped completely with the center of the bigger cluster associated with extension strength.

#### Structural brain damage associated with specific aspects of finger-dexterity loss

VLSM conjunction analysis^14, 17, 18^ (Fig. 4A; Table 3) was done to explore which brain structures were specifically associated with finger-individuation in one direction but not the other, as well as brain structures associated with finger-individuation in both directions. To be able to compare lesion effects on individuated flexion and individuated extension, only voxels passing a z-score threshold of 2.32 (corresponding to p < 0.01) were considered significant. Most such voxels were associated with finger individuation in both flexion and extension directions (60% of the voxels, n = 322), and were grouped in a single cluster which involved SLF-II (32%), CRP (16%), M1 (15%), STR (14%), M1-CST (9%) and PM-CST (5%). Damage to about 35% (n = 188) of the significant voxels affected exclusively individuation of finger movement in the flexion direction. These flexion-selective voxels were arranged in three clusters. The largest and most significant one (151 voxels, zmax = 3.61, MNI coordinates, -32 –20 54; see Fig. 4A) involved mainly white-matter projection and association tracts: M1-CST (38%), SLF-II (31%), PM-CST (13%), CRP (12%), STR (12%), S1-CST (9%) and M1 (6%).

**Figure 4.**
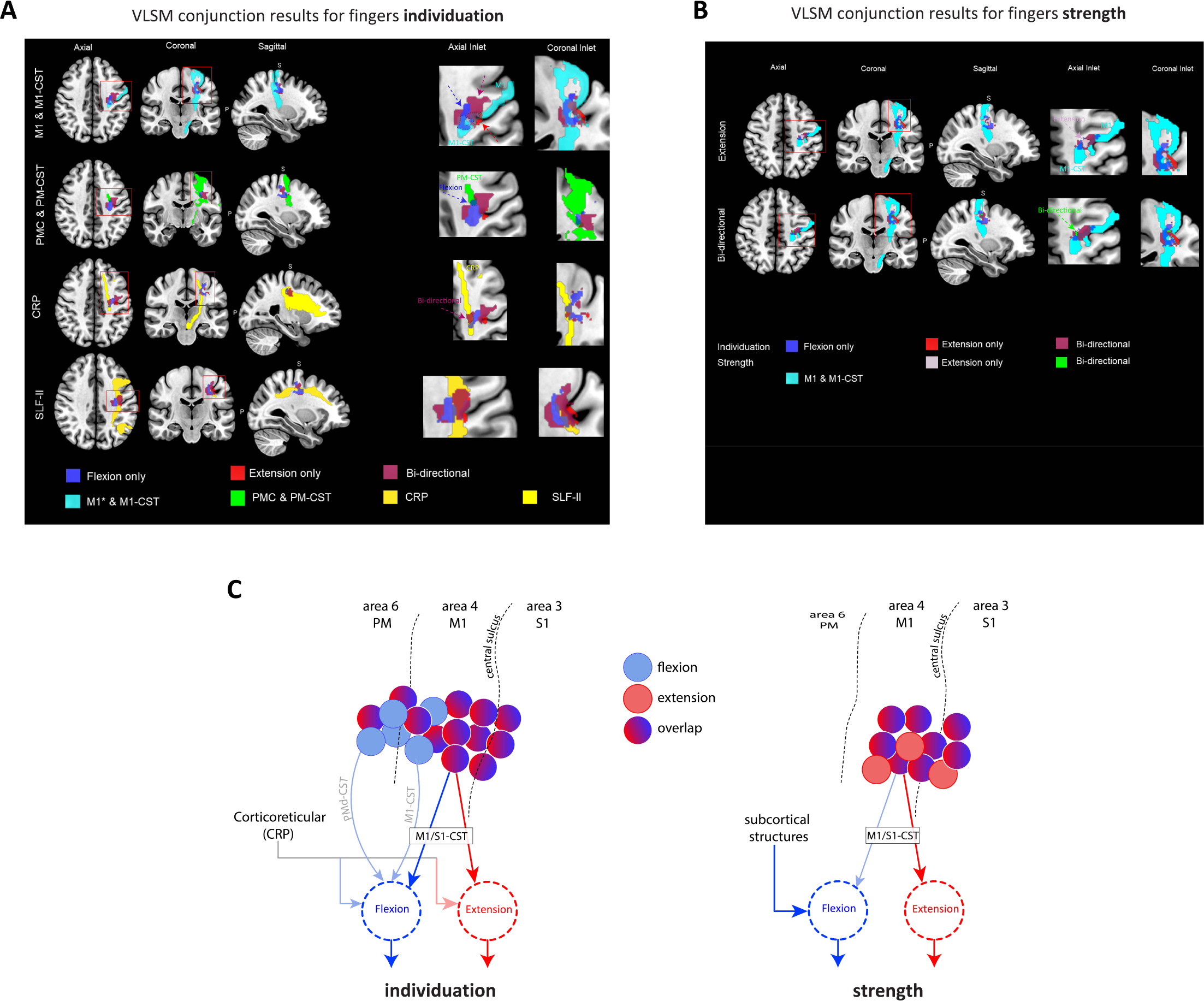
VLSM conjunction analysis. (A) VLSM conjunction analysis depicting voxels in which the existence of damage was associated with significant impairment of finger individuation capacity in flexion only (blue), extension only (red), or both (bi-directional, purple) and the major brain areas in which these voxels were detected. The depicted voxels survived a lenient criterion of p < 0.01 (Z > 2.32). Brain structures: M1 = primary motor cortex (without lower-limb and trunk); PMC = premotor cortex; CST-M1, CST-PMC = corticospinal tract fibers originating in M1 and PMC, respectively; CRP = cortico-reticulospinal pathway; SLF-II = 2^nd^ division of the superior longitudinal fasciculus. (B) VLSM results for finger strength overlayed on VLSM conjunction results for individuated fingers movement, together with primary motor cortex and M1-CST (individuation results are the same as the first row of Figure 4A). Only voxels associated selectively with extension (pale pink) or bi-directional (green) finger-strength are depicted since no voxels were associated selectively with flexion-strength. The depicted voxels survived lenient criterion of p < 0.01 (Z > 2.32). Brain structures: M1 = primary motor cortex (without lower-limb and trunk); CST-M1 = corticospinal tract originating in M1. (C) Contribution of cortico-reticular and corticospinal fibers originating in different areas of the sensory-motor and premotor cortices to finger individuation capacity and strength, in flexion and extension.

**Table 3.**
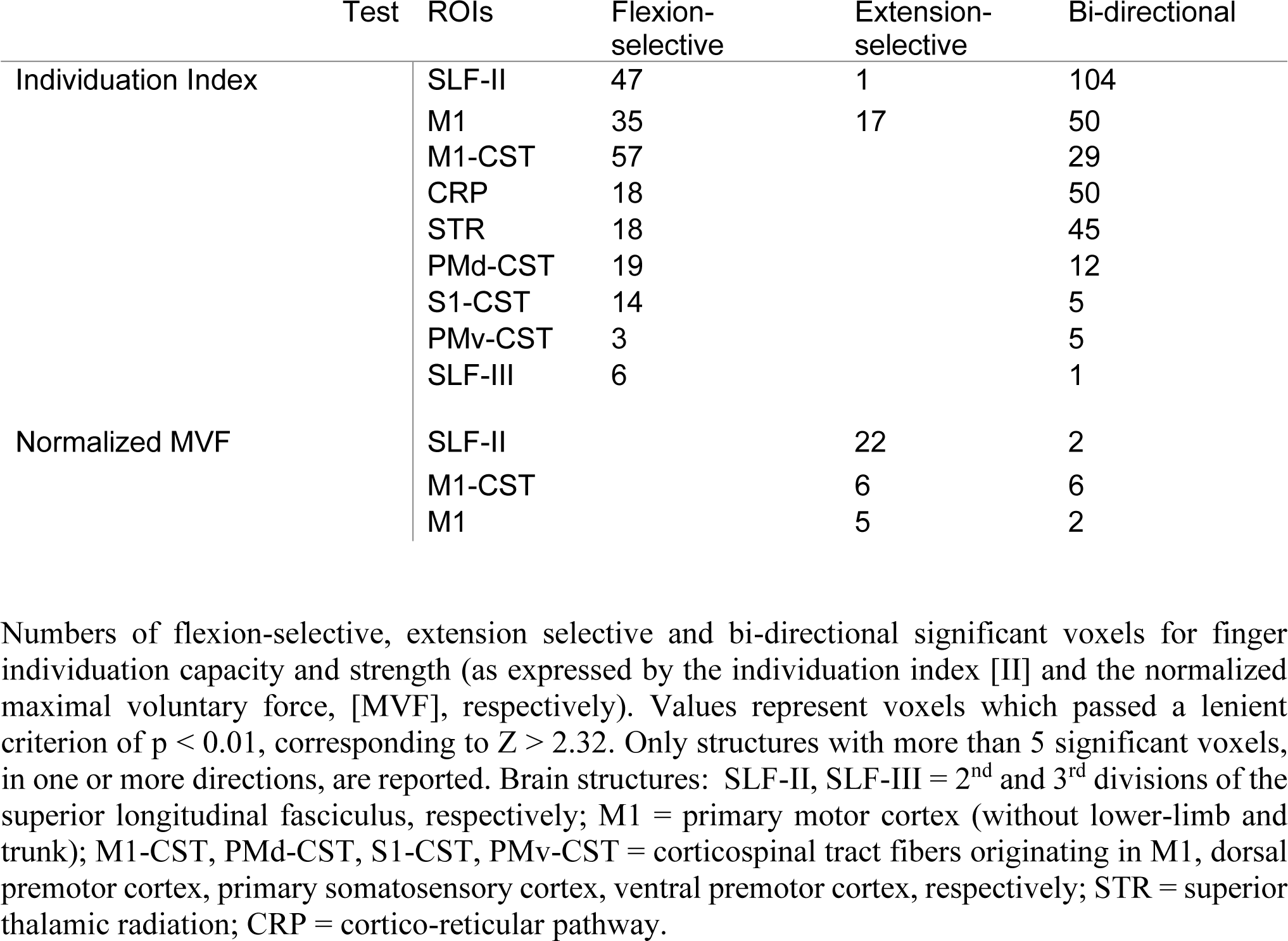
VLSM conjunction analysis. Numbers of flexion-selective, extension selective and bi-directional significant voxels for finger individuation capacity and strength (as expressed by the individuation index [II] and the normalized maximal voluntary force, [MVF], respectively). Values represent voxels which passed a lenient criterion of p < 0.01, corresponding to Z > 2.32. Only structures with more than 5 significant voxels, in one or more directions, are reported. Brain structures: SLF-II, SLF-III = 2^nd^ and 3^rd^ divisions of the superior longitudinal fasciculus, respectively; M1 = primary motor cortex (without lower-limb and trunk); M1-CST, PMd-CST, S1-CST, PMv-CST = corticospinal tract fibers originating in M1, dorsal premotor cortex, primary somatosensory cortex, ventral premotor cortex, respectively; STR = superior thalamic radiation; CRP = cortico-reticular pathway.

| Test | ROIs | Flexion-selective | Extension-selective | Bi-directional |
| --- | --- | --- | --- | --- |
| Individuation Index | SLF-II | 47 | 1 | 104 |
|  | M1 | 35 | 17 | 50 |
|  | M1-CST | 57 |  | 29 |
|  | CRP | 18 |  | 50 |
|  | STR | 18 |  | 45 |
|  | PMd-CST | 19 |  | 12 |
|  | S1-CST | 14 |  | 5 |
|  | PMv-CST | 3 |  | 5 |
|  | SLF-III | 6 |  | 1 |
| Normalized MVF | SLF-II |  | 22 | 2 |
|  | M1-CST |  | 6 | 6 |
|  | M1 |  | 5 | 2 |

The other two clusters were much smaller and less significant: cluster 1 (11 voxels, zmax = 2.37, MNI coordinates –44 –4 28); cluster 2 (26 voxels, zmax = 2.59, MNI coordinates, –38 –12 44) and involved mostly voxels within the primary motor cortex (M1). There was a single small extension-selective cluster (23 voxels, zmax = 3.26, MNI coordinates, -26 –16 48; see Fig. 4A) involving mainly M1 (74%) and S1 (17%). Finally, a conjunction analysis was carried out to explore brain voxels in which the existence of damage affected exclusively finger individuation, exclusively finger strength, or both individuation and strength. We found out that 79% of the voxels associated with finger strength (either flexion or extension) were also associated with bi-directional finger individuation (n = 52 voxels; M1-CST 12%, M1 10%; Fig. 4B, Table 3). The number of voxels associated with both finger strength and selective fingers individuation was scarce (4 voxels for individuated flexion, no voxels for individuated extension).

In summary, we have found voxel clusters involving the primary motor cortex (M1) and white-matter tracts associated with motor control (M1-CST, PM-CST, CRP, SLF-II, STR) where the existence of damage affected finger individuation capacity only in flexion or in both flexion and extension directions. Voxel clusters associated exclusively with individuated extension were scarce and localized at the junction between M1 and S1 (involving both). Voxels associated with finger strength (both directions) were largely associated also with bi-directional finger individuation and involved M1 and its corticospinal relays (M1-CST). Interestingly, voxels associated with individuation capacity in one direction only, formed 35% (flexion selective) and 5% (extension selective) of the total number of voxels in which the existence of damage affected finger individuation. In contrast, voxels associated with force generation in only one direction of finger movement, formed 0% (flexion selective) and 85% (extension selective) of the total number of voxels in which the existence of damage affected finger strength.

## Discussion

In this study, behavioral analyses of finger individuation and force generation disclosed marked direction-dependent differences. Lesion-symptom relationship analysis at the voxel level completed the picture by pointing to distinct neural substrates subserving these elements of motor control which are crucial for dexterous performance of daily tasks. Behavioral characterization provided clear evidence for an advantage - both in fine and gross motor control (individuation, strength) - when finger movements are executed in the flexion direction compared to the extension direction. We could also show that the increased precision of control in flexion does not stem from properties the human hand’s anatomy, but rather reflect distinctions in the underlying neural mechanisms that control the movements in these directions. Voxel-based lesion-symptom mapping identified causal relationships between finger individuation capacity and restricted regions within M1, S1 and corticospinal fibers descending from these regions and from the premotor cortex, as well as other white-matter fiber tracts (SLF, STR and CRP). Although there was a great deal of anatomical overlap between regions implicated in individuation capacity in flexion and extension, we were able to identify cortical and white-matter structures that were associated exclusively with individuated finger flexion and to a lesser extent, exclusively with individuated finger extension (the latter, unlike the former, did not involve long white-matter tracts but only M1 and S1 regions). A small part of the region implicated in both individuated flexion and extension, near the place of origin of M1-CST fibers, was implicated also in force generation.

### Neural control, not mechanical coupling, underlies the behavioral differences between finger flexion and extension

Traditionally, finger individuation capacity is considered a crucial component of manual dexterity. This capacity is thought to reflect an aspect of motor control which is different from other characteristics of dexterous manual activity, such as precision grip and finger strength^19, 20^. Early studies in macaque monkeys^21^ showed that almost all movements of instructed digits were accompanied by motion of the other digits. Thumb flexion was accompanied by relatively little concomitant motion of other digits, and the monkeys’ capacity to individuate finger movements in extension was inferior in comparison to movement in flexion. Yet, flexion/extension differences in different fingers were sparse and inconsistent and finger strength was left unexplored in this study ^21^. Finger individuation in humans was studied in *isotonic* and *isometric* conditions. In the former condition, movement was usually examined against minimal or no external load. It was shown that while intending to move only a single digit, humans often move the other digits as well. Self-paced oscillatory flexion and extension movements of a single digit were accompanied by notable movements of the other digits, and more so following brain damage^2, 22^. Flexion and extension finger movements of humans were found to be less individuated for the middle and ring fingers compared to the thumb, index and little fingers. Individuation indexes for human flexion and extension movements were noticeably higher than the corresponding values acquired from macaque monkeys. In the *isometric* condition, individuation capacity is studied in terms of subjects’ tendency to unintentionally generate force (*enslaving*) in uninstructed digits while intending to exert isometric force in a single instructed digit or in a specified combination of instructed digits^10, 11^. Individuation and enslaving may be seen as inversed phenomena (lower individuation capacity corresponds to a higher degree of enslaving). The advantage of the isometric task is that it allows for quantifying individuation capacity in graded strength requirements.

Our results are consistent with earlier reports showing that enslaving is lower during force generation in the instructed finger in flexion compared to extension^23, 24^. We add novelty to existing knowledge by characterizing human individuation capacity, not only in the single instructed digit condition, but also in more complex conditions, where individuation is required from multiple combinations of instructed digits, as it occurs in natural situations demanding manual dexterity, like guitar or piano playing. We demonstrate that differences between flexion and extension in the magnitude of enslaving appear also in conditions of multi-digit movement, beyond specific enslaving patterns and graded force-emission requirements from the instructed digit. We attempted here (for the first time, as far as we know) to isolate the contribution of mechanical coupling, due to properties of the hand’s anatomy, to the difference in the magnitude of enslaving seen in flexion and extension. We show (in experiment 2) that when a digit is moved passively by the robot, while measuring isometric forces generated unintentionally in the unmoved fingers, mechanical coupling prevents the instructed digit from being moved by the robot in isolation. This finding corroborates previous reports^25^. The novelty here is in showing that mechanical coupling is significantly higher in passive flexion compared to passive extension. This finding rules out the possibility that mechanical coupling explains why active flexion is accompanied by less enslaving (superior individuation capacity) compared to active extension. It rather suggests that individuation advantage in active flexion compared to extension stems from dissimilar neural control of finger movement in the two directions. Note that multi-site surface EMG recording during the passive stretching confirmed that forces generated in unmoved digits were not caused by neurally-mediated muscle contractions, thus ruling out possible spinal-cord responses to either muscle stretch or focal discomfort/pain as mediators of force generation^26, 27^. This leaves inter-digit mechanical coupling of soft tissues, skin and ligaments as the likely source of force generation in the unmoved digits during passive motion of the instructed digit. Further study is needed to quantify the contribution of these factors to limiting the individuation capacity in active finger movement.

### Neural correlates of individuated finger movement in flexion and extension

Dexterous execution of hand movements to achieve desired behavioral goals depends on integrated neural inputs received from various sources within the central nervous system, e.g., spinal segmental interneurons^28–30^, propriospinal neurons^3, 31^, neurons in the pontomedullary reticular formation^32–34^, the red nucleus^35, 36^, and the cerebral cortex^20, 37–39^. Inputs received from these and other sources directly, and more often indirectly, modulate the activity of α-motoneurons, which emit the final neural commands for contraction of muscles and production of movement. With the evolution of the 6-layered neocortex in mammals, cortical involvement in motor control increased, and the CST, having diverge origins in sensory-motor, premotor, and cingulate cortical regions, became a principal mediator of supra-spinal motor control^40, 41^. The emergence, later in evolution, of direct cortico-motoneuronal (CM) projections, reaching peak development in the more advanced primates, especially in humans, was associated with deeper involvement of the cerebral cortex in limb motor control. Such involvement became a necessity when fore limbs became engaged in complex cognition-loaded tasks, like tool manufacturing and use, rather than in locomotion ^42, 43^. Most CM projections arise from ‘new’ M1 (Brodmann’s area 4p, buried in the central sulcus) and from adjacent area 3a, and terminate in the spinal ventral horn, close to cell-bodies of α-motoneurons^41, 44^. Research in non-human primates showed that CM projections from ‘new’ M1 cells reach spinal motoneuron pools innervating multiple muscles^45^. Recording of activity from single neurons in M1 hand-knob region, during individuated flexion/extension movements in single digits or the wrist, disclosed considerable regional overlap in M1^5^, and firing in single CM neurons preceding a variety of individuated digit movements^6, 7^. Along the mammalian evolutionary scale, the proximity of CST terminations to α-motoneurons correlates roughly with the emergence of dexterous performance of more and more complex tasks^46^. Deficits that result from damage to M1 or the CST as well as functional neuroimaging research^4^ attest to the crucial importance of this system in generating individuated finger movements. In accord, our VLSM analysis in first-ever stroke patients, disclosed brain voxels within M1 and M1-CST in which the existence of damage played a causative role in manifestation of impaired finger-individuation, both in flexion and extension.

These however were not the sole regions where damage by stroke affected finger individuation. Damage to voxels within other association (SLF) and projection fiber tracts (PMd-CST, STR, CRP, S1-CST) proved to be significant as well (Table 2; Fig. 4A), thus showing that supraspinal control of individuated finger movement incorporates inputs received from divergent sources in the human cerebral cortex. This finding is in accord with animal research^4^ showing convergence of inputs from various cortical sites when a single muscle is activated and overlap of cortical circuits involved in activation of different muscles^47, 48^. Research in non-human primates employing intracortical micro-stimulation to elicit digit movements, informed on functional connectivity patterns between digit representations in M1 and both ventral and dorsal cell aggregates in the premotor cortex, with varying degrees of anatomical segregation and overlap in the latter for different movements^49, 50^. Of note, damage no less than 151 brain voxels within the second division of the SLF affected the capacity for finger individuation in flexion, and damage to 105 brain voxels in the trajectory of this fiber tract, affected finger-individuation capacity in the extension direction. In both movement directions, the numbers of voxels associated with finger individuation (and the percentage of the brain structure they occupied) was larger for SLF-II than for all other cortical and sub-cortical brain regions (Table 2, Fig. 4A), attesting for the importance of robust fronto-parietal connectivity in dexterous task-driven and attention-demanding hand movement, as reflected by the capacity to individuate finger movements in our present research tasks.^51–53^

### Flexion-selective brain voxels in the motor network

We have shown that finger flexion is accompanied by less unwanted enslaving of other fingers compared to finger extension, despite the fact that flexion movement induces more mechanical coupling than extension. This finding points to differences in the neural control of finger movements in flexion and extension. VLSM conjunction analysis sheds new light on this finding by showing that a much larger number of brain voxels, in different parts of the motor network, relate to individuation capacity in the flexion direction compared to the extension direction. While large numbers of brain voxels located in sensorimotor, premotor cortices and in white matter association and projection fiber tracts, revealed causative relationship with finger individuation capacity both in flexion and extension, we found substantial numbers of other brain voxels, in the same regions, that revealed a selective association with finger individuation capacity only in flexion (Table 3, Fig 4A). In sharp contrast to this finding, selective relationship with individuation capacity in extension was revealed by only a meager number of M1 voxels. The large majority of flexion-selective voxels were found in sensory-motor and premotor cortices and in the trajectories of cortico-spinal and cortico-reticular fibers descending from these regions. A sizable voxel cluster involved also the second division of the SLF. The much larger representation of flexion-selective, compared to extension-selective, neurons in the motor network explains the superiority of finger individuation capacity in the flexion direction. It may also be related to the phenomenon, often observed in stroke survivors, where the residual capacity for grasping and holding objects is higher than the capacity to afterwards carefully release the objects.^40, 54–56^. The largest number of brain voxels showing a selective association with finger individuation capacity in the flexion direction, was found to reside in M1-CST (cortico-spinal fibers originating from the primary motor cortex). This finding is in accord with a recent DTI lesion study of finger strength vs. control after stroke, which implicated the M1-CST in individuated finger flexion^57^. Our finding of premotor contribution to individuation capacity of finger movement in flexion, is in accord with studies in non-human primates implicating propriospinal neuronal (PN) relays of premotor-CST fibers, in finger individuation and dexterous hand movement, especially after selective CST lesioning^3, 58–60^. Up-regulation of projection fibers originating in the premotor cortex and acting on cervical propriospinal neurons was shown in humans after stroke^61, 62^. Thus, the premotor contribution to dexterous finger movement post stroke, shown in the current study, is likely to be mediated by propriospinal neurons in cervical segments of the spinal cord. Note that a considerable number of brain voxels in primary motor and premotor cortices, in which the existence of damage affected finger individuation capacity, did not show a directional selectivity and were associated with both finger flexion and extension. However, input from a specific group of cells within these regions is primarily biased towards flexion movement and adds more degrees of controllability to finger individuation in the flexion direction.

### Different neural mediation of finger strength and individuation

Like individuation capacity, the ability to generate graded levels of force at will is a crucial pre-requisite for dexterous performance of daily tasks. When lifting an object, the grip force one generates needs to be adapted to the object’s weight (such that it doesn’t fall down) and consistency (such that it doesn’t break). The classical studies of Lawrence and Kuypers in non-human primates^1, 63^ showed that individuation and strength, two major aspects of hand dexterity, are not controlled by one and the same neural mechanism. While individuation depends mainly on the integrity of cortico-spinal projections originating from M1, strength seems to receive contributions from additional sources, such as the reticulo-spinal system. Our VLSM findings support the notion of at least partly separated control systems for finger individuation and force generation, by showing different profiles of significant brain voxels for the two. Note however that contrary to what could be expected, large numbers of brain voxels (not only within M1 and M1-CST, but also in the premotor cortex and in the trajectory of cortical projections to the reticular system), were found to be associated with finger individuation, whereas much fewer voxels, in M1 and SLF-II, were associated with finger strength (Table 3). The detection of relatively few anatomical correlates of finger force in the current analysis, may be methodological constrains of VLSM, that cannot detect lesion-symptom relationship outside the minimal overlap region, which included only regions within the core of MCA vascular territory that included the CST but only small part of the CRP. Additionally, CST descends in a much smaller area relative to the CRP, thus is far more vulnerable to focal damage^64, 65^ (compare the CST and CRP on the first and third rows, respectively). The partial dissociation of neural mechanisms controlling finger individuation and strength (depicted in Figures 3F and 4B), is reflected also in the inverse anatomical patterns of individuation and strength in flexion and extension (much more flexion-selective voxels in VLSM analysis of individuation capacity, in contrast to the existence of extension-selective but no flexion-selective brain voxels in the VLSM analysis of finger strength; Table 4). Our findings, pointing to a difference between the neural control of finger individuation and finger strength, are in accord with a recent study that disclosed different patterns of recovery for finger strength and finger individuation in the first year after the onset of stroke^57^.

### The impact of damage to fiber tracts

The mapping of lesion-symptom relationships in the current study is driven from a cohort of patients who survived a stroke of mid-range severity, giving rise to substantial involvement of subcortical structures and white-matter tracts in core middle-cerebral-artery territory, but leaving large areas of the cerebral cortex without sufficient lesion overlap to permit accurate assessment of lesion-effects there (Figure 3A; see *Limitations of the study*). With respect to fiber-tract damage, we note the specific importance of a region (corresponding in the left to MNI coordinates [–26, –18, 40]), traversed by projection fibers originating both in M1 and PMC, in which the existence of damage was associated with selective decrement of finger individuation capacity in the flexion direction. Damage to nearby voxels (within the same fiber tracts) affected finger individuation both in flexion and extension. Damage to a much smaller number of brain voxels in M1 was associated with selective decrement of finger individuation capacity in the extension direction. These findings go beyond the current understanding of neural mechanisms involved in manual dexterity, by pointing to the existence of two separable neuronal populations in primary motor and premotor cortices – one involved in bi-directional finger individuation and another involved selectively in individuation of finger movement in flexion. The existence of an extensive portion of grey and white matter, within the motor system, which is involved in the control of finger movements in flexion, and the overwhelming lack of a comparable substrate devoted to fine control of finger movement in extension, explain why behavioral testing in healthy subjects reveals a significant superiority of individuation capacity in finger flexion compared to finger extension. Of note is the fact that flexion-selective control was not found for the other major determinant of manual dexterity – the capacity to generate force in the fingers. Here actually the analysis of lesion effects on normalized MVF pointed to a relatively small number of brain voxels, within the trajectory of white-matter association and projection tracts (SLF-II, M1-CST) and within M1, that were involved in mediation of force generation in finger extension, with no parallel brain substrate devoted exclusively to force generation in the finger flexion movements. Figures 4A and 4B depict the distinctive effects of lesions to distinct white matter tracts and their cortical sites of origin. Figure 4C summarizes in a schematic form the above findings.

### Limitations of the study

To the best of our knowledge, the current study is the first attempt to explore the neural mechanisms underlying finger individuation capacity in different movement directions in humans. The paradigm we used for that aim has several limitations that should be noted. First, patients were recruited during the early subacute phase after stroke, while receiving in-patient rehabilitation. This means that the structure-function relationships revealed by our VLSM analysis reflect not entirely the brain’s normal physiology but also its modulation by natural and treatment-related plastic changes^66, 67^. Second, given our aim to assess comparatively finger flexion and extension, we recruited patients that were able to undergo the assessment of individuation capacity in the robot at both movement directions. This requirement dictated exclusion of stroke patients with little or no finger movement, as well as patients with no apparent motor deficit (as determined by clinical assessment, and patients’ self-reports). This approach may have induced a selection bias; hence our results may not apply for stroke patients with too severe or too mild paresis. Third, there are technical limitations to our setup: the computed individuation index derives from an isometric task, which differs from the individuation required in natural tasks like piano playing. Also, our apparatus was limited to a maximal force measurement of 20 N, thus subjects (some of them with greater finger strength) were instructed not to exceed this level. Such requirement may have induced a potential ceiling effect to our results, similarly to the one seen in many clinical assessments^68^ . Finally, VLSM analysis has several known limitations ^69^. To ensure sufficient statistical power, only brain voxels damaged in >15% of the patients were entered into analysis, resulting in reduced coverage of brain regions that are infrequently lesioned (see Fig. 3A for the brain territory that entered analysis). As the more dorsal regions in the lateral hemispheric surface are often spared in MCA stroke, upper-limb paresis frequently stems from CST damage in corona-radiata or internal-capsule level, with the hand knob in M1 remaining intact or only partially involved. This often leads to under-estimation, in VLSM studies, of M1 importance to upper-limb function. Similar under-estimation of importance may apply for adjacent cortical regions, like the hand area in S1 and the dorsal premotor cortex. Given this limitation we focused on damage to white-matter tracts for whom the place of origin in the cortex is known.

In summary, our data support the inclusive view that the neural substrate underlying individuated finger movement in flexion and extension overlap remarkably in M1, PMC and their related projections. In addition to this common neural substrate, individuated finger movement in the flexion direction depends on the integrity of more extended, flexion-specific, areas within the same brain structures. This extended cerebral representation of individuated movement in flexion compared to extension is likely to explain the superior individuation capacity of finger movement in flexion, that is revealed in behavioral testing. The other major determinant of manual dexterity – finger strength – seems to have a considerably different functional anatomy.

## Materials and Methods

### Participants

A total of 83 participants were recruited for this study. **In experiment 1**, a cohort of 13 right-handed young healthy participants (8 females; age, mean±STD: 28.2±2.9 years) was recruited and given monetary compensation for a two-hour session (100 ₪≈ $30). One participant was dropped after the first hour and consequently given half the compensation. **In experiment 2**, a new cohort of 20 right-handed young healthy participants (11 females; age 31.7±6.1 years) was recruited and given monetary compensation for a one-hour session. Both experiments were approved by the institutional review board and ethics committee of the Technion - Israel Institute of Technology. To confirm that enslaving forces measured in the passive condition in Exp. 2 reflect mechanical coupling between the digits and are not neurally-mediated contractions in the context of passive stretching, an additional 7 right-handed healthy control participants (3 females, age 36.3±7.9 years) were recruited and performed experiment 2 while we recorded EMG activity from finger/hand flexor and extensor muscles. **In experiment 3**, we recruited a cohort of 32 hemiparetic stroke patients undergoing in-patient rehabilitation (9 females; age 60.5±10.6 years; time after onset 40.9±17.0 days, see Table 1 for patients’ details). Patients were recruited for the study if they met the following inclusion criteria: (1) age: 20–80 years; (2) right hand dominance; (3) first-ever unilateral ischemic or hemorrhagic stroke; (4) time after stroke onset 14-60 days; (5) upper-limb paresis with residual force-generation capacity of at least 0.5N in all digits in at least one direction (flexion/extension); (6) cognition and language allowing provision of informed consent and following task instructions. We excluded patients with one or more of the following: (1) unstable medical status; (2) prior neurological/psychiatric disorders; (3) prior orthopedic/rheumatologic disorders affecting upper-limb function; (4) cerebellar involvement; (5) sensory problems that may prevent reporting pain during interaction with the robot; (6) skin breakdown or wounds in places where the hand contacts the robot; (7) participation in another interventional study for upper limb rehabilitation. For detailed stroke patients’ characteristics, see Table 1. For comparison, we recruited a cohort of 11 healthy age-matched controls (6 females; age 56.3±10.0 years). There was no significant difference in age between the patient and control populations (𝑡_41_ = 1.1439, 𝑝 = 0.2593). Experiment 3 is part of a pre-registered clinical trial (NCT04229329) that was approved by the Helsinki committee of the Loewenstein Rehabilitation Medical Center. All patients signed an informed consent.

### Overview of finger dexterity experiments

#### Apparatus and user interface

In all our experiments, participants (healthy and stroke patients) were seated facing a computer screen with the right hand (or paretic hand in patients) placed in a neutral posture on a dexterity robot (Amadeo, Tyromotion®) (**Fig. 1A**). The robot is built of five mini handles to which each digit can be connected using magnetic thimbles wrapped around the fingertips with band-aid tapes. Personal physical adjustments were made to ensure comfort and suitability to the participant’s hand dimensions. The handles of the robot can work in two modes: isotonic or isometric. In the *isotonic* mode, the handles can be moved freely in the flexion-extension axis and positions can be measured between the two mechanical end stops with 0 representing the maximal extension and – 1 representing the maximal flexion. In practice, the movement of each digit was restricted within the participant’s range of motion (ROM). The ROM was personally configured by the fully opened and fully closed positions each participant was able to actively reach. In the *isometric* mode, the handles are locked in a pre-defined position (configured once in the beginning and remaining constant through the experiment) and forces are measured in the flexion/extension directions. While in healthy participants, handle position was defined individually as the middle of the ROM, in stroke patients, it was defined manually by a certified physical therapist. In this manual positioning, patients were seated on a chair with a back support in the following sitting posture: 90 degrees of the seat back angle, pronated forearm, ∼20 degrees extension of the elbow, fingers in a start opening position resembling grasping a tennis ball.

The dynamic range of measured forces was ±20 N, with negative values indicating force in the flexion direction and positive values indicating force in the extension direction. To precisely measure the forces during isometric activity while considering baseline forces produced by the hand’s posture and due to the digits’ weights, we subtracted average forces measured by the sensors during an approximately 2-second time period in which participants were instructed to be at rest. This zeroing procedure was performed once in the beginning of the experiment (once in the beginning of each block for patients). Analog force and position signals were digitized and sampled internally at 60 Hz. Timestamps were added to the samples, and were integrated with a customized MATLAB script (The MathWorks, Inc., Natick, MA) enabling real-time measurements, presentation, and analysis. Using Psychtoolbox-3, live visual stimuli were presented on a computer screen and guided the participants through the tasks. Following a short demonstration of the isotonic and isometric modes and an explanation of the graphical user interface (GUI), participants were provided with an overview and detailed description of the tasks.

#### Experimental protocols

The aim of ***experiment 1*** was to characterize individuation and force-generation capacities of healthy participants during finger flexion and extension. Participants performed three isometric tasks with their right dominant hand: a finger-strength task, a single-finger individuation task and a multi-finger individuation task (**Fig. 1B**). The working position for all three tasks was defined as the middle of the ROM. Our objective in ***experiment 2*** was to quantify the extent to which mechanical coupling between fingers contributed to movement of uninstructed fingers as observed in healthy participants. Participants underwent a passive stretching task by the robot and performed the finger-strength and the single-finger individuation tasks using their right dominant hand, in a counter-balanced order (**Fig. 2B**). Whereas the working position for the finger-strength task and the single-finger individuation task was defined as before, the passive stretching task required two different starting positions depending on the direction of movement (flexion/extension began from 90% of the fully opened/fully closed positions, respectively). In ***experiment 3*** we sought to study the effects of brain lesion topography on finger individuation and force-generation capacities in flexion vs. extension movement directions. Sub-acute stroke patients and age-matched healthy controls performed the finger-strength task and the single-finger individuation task. Stroke patients performed the finger-strength task on both the paretic and intact hands. The single-finger individuation task was done only with the affected hand, in two separate sessions (1-3 days apart) to obtain an averaged measure of performance, in view of fluctuations commonly seen among sub-acute stroke patients. Healthy controls were evaluated on both their right (dominant) and left hands in a counter-balanced order (**Fig. 3B**). During this experiment, the working position was defined manually by a certified physical therapist (see above for the description of the manual positioning).

#### Assessment of finger strength

Prior to evaluating the individuation aspect of finger dexterity, we measured the maximal voluntary contraction force (MVF) of each finger in flexion and extension directions. Each participant performed 20 trials (5 digits x 2 directions x 2 repetitions) in this task. In each trial, participants were asked to produce as much force as they could in a specific digit and direction and maintain that force for 3 seconds (2 seconds for patients). Here, participants were not limited to using only the instructed digit to produce the maximal force but were allowed to produce forces also in the rest of the digits. The trials were executed in a fixed order: flexion than extension, starting with the thumb and ending with the little finger. The maximum force in each trial was calculated as the 95th percentile of the instructed digit’s absolute force data. Each action was repeated twice. If the ratio between the forces generated in the two repetitions (smaller value divided by the larger value) was below 60%, the higher value was considered as the MVF, otherwise the average value was considered as the MVF. Due to limitation of the robot sensors, participants were allowed to generate forces only within the dynamic range (±20 N). In case the force exceeded this value, a warning signal appeared, and participants were told to slightly reduce force. Out-of-range forces were automatically floored to ±20 N.

#### Single-finger individuation task

The capacity to move an instructed finger, without moving the other fingers at the same time, is a basic pre-requisite for dexterous performance of various daily manual tasks. To quantitatively assess this capacity, participants were instructed to actively exert force, each time with a different single digit, in a specified direction (flexion, extension), and maintain the generated force in one of four target levels (20, 40, 60 or 80% of the MVF), while keeping all other digits as static as possible. Each participant performed 120 trials (5 digits x 2 directions x 4 force levels x 3 repetitions) in this task. The trials were executed in a fixed order: flexion and extension directions were alternated, force levels were monotonically increased from 20 to 80% of the MVF level, starting with the thumb and ending with the little finger. In each trial, participants had to generate in the instructed finger and direction (flexion, extension) an amount of force that brings a horizontal white line into the middle of a target green rectangle on the screen, and maintain that force (i.e., maintain the line within the rectangle on the screen) for 2 seconds, with allowed force deviations from target of 10% (15% for patients). At the same time, participants were instructed to see that the other digits stay at rest (visual guidance was provided by gray rectangles on the screen depicting forces generated in the uninstructed digits, with tolerance of 20% around zero N). (**Fig. 1C**). Participants had to complete each trial within 5 seconds (7 seconds for patients), after which the trial was terminated. The next trial started after a short rest (inter-trial interval [ITI] of 2 and 4 seconds, for healthy controls and stroke patients, respectively). Trials with missing or invalid data from at least one force sensor were excluded. In total, 0.5% of the trials in experiment 1 were excluded, 1.2% in experiment 2, 2.7% of the trials in the stroke group and 0.2% for the healthy controls in experiment 3.

#### Multi-finger individuation task

Here, individuation capacity was examined in a more complex task, demanding multi-digit individuated contraction (“chords”). In each trial, participants were instructed to perform a chord (out of 25 possibilities, 10 x two-digit, 10 x three-digit and 5 x four-digit) either in the flexion or extension direction, and the required force was normalized across digits (i.e., 25% of MVF calculated separately for each digit and direction).

Each participant performed 150 trials (25 combinations x 2 directions x 3 repetitions) in this task. The trials were executed in a fixed order – flexion and extension directions were alternated, starting from 2-digit chords, proceeding to 3-digit chords and ending with 4-digit chords. In each chord, participants had to generate the specified force in the instructed fingers, maintaining for 2 seconds horizontal white lines on the screen in the middle of green rectangles indicating the target forces with allowed force deviations from target of 10%. At that time, participants were instructed to see that the other digits stay at rest (visual guidance was provided by gray rectangles on the screen depicting forces generated in the uninstructed digits, with tolerance of 20% around zero N) (**Fig. 1C**). Participants had to complete each trial within 5 seconds (7 seconds for patients), after which the trial was terminated. The next trial started after a short rest (inter-trial interval [ITI] of 2 seconds).

#### Assessment of inter-digit mechanical coupling

The aim here was to quantify, in healthy subjects, the extent to which mechanical coupling between the fingers contributes to movement of the uninstructed fingers in the individuation tasks. Participants were instructed to maintain all fingers at rest and refrain from any voluntary contraction while each digit in its turn was moved passively by the robot. In particular, the instruction regarding the passively moved digit was to neither resist the robot nor assist it. The other un-moving digits were locked in their positions and forces generated in them were measured. The task was divided into a *ROM* and *Velocity* blocks. In each trial of the *ROM* block, the robot moved a single digit through one of four distances (20, 40, 60 and 80% of ROM) at the maximally allowed robot velocity (0.2 m/sec). In each trial of the *Velocity* block, the robot moved a single digit through a fixed distance (set at 8% of the ROM) in one of three velocities (0.05, 0.1 and 0.15 m/s). In total, each participant performed 120 trials (5 digits x 2 directions x 4 distances x 3 repetitions) in the *ROM* block and 90 trials (5 digits x 2 directions x 3 velocities x 3 repetitions) in the *Velocity* block (1.2% of all the trials were manually excluded). In both blocks, the trials were executed in a fixed order: flexion and extension directions were alternated, distances/velocities were monotonically increased, and digits were passively moved by the robot, starting with the thumb and ending with the little finger.

#### EMG

Electromyography (EMG) signals were recorded both during active and passive tasks using a surface EMG system (Delsys Trigno™) from 6 superficial muscles normally activated during finger flexion and extension: flexor pollicis brevis (FPB), flexor digiti minimi brevis (FDM), flexor digitorum superficialis (FDS), first dorsal interosseous (FDI), abductor pollicis longus (APL) extensor digitorum (ED). Muscle activity was recorded using 6 high-density Ag/AgCl electrodes in a belly-tendon montage while the participant performed the tasks. The signal from each electrode was sampled at 1,000 Hz, demeaned, rectified and low-pass filtered (fourth order butterworth filter, fc = 40 Hz). Finally, the processed EMG signals were averaged across each trial over a 3.5-s time window starting from the time when the instructed finger(s) first moved (in active condition) or when the robot moves the finger (in passive condition).

### Data analysis

#### Preprocessing of kinetic data

Force data was pre-processed offline using custom scripts written in MATLAB (The MathWorks, Inc., Natick, MA). We applied the following processing steps: (1) low-pass filtering, the raw force data of each sensor was filtered using a gaussian filter (σ = 5) to eliminate the electrical noise from the sensors and smooth the force traces. (2) baseline-correction, the forces at the beginning of each trial (baseline forces) were calculated over the initial 1.5 seconds of the trial in three segments (0-0.5, 0.5-1.0, 1.0-1.5 seconds). In order to verify that the finger was at rest, the standard deviation of each segment was calculated and the segment with the lowest deviation was chosen. Then, the mean force of that segment was calculated and set as the baseline force of a given digit. The overall baseline forces were subtracted from the data, resulting in modified forces which start at 0 N for each trial. (3) forces onset and offset, those timestamps were marked as the first and last times the instructed digit’s force passed 50% of its peak (minimum for flexion, maximum for extension), respectively. Since multi-finger individuation task involves more than one instructed digit, the longest range was selected in this case (minimum over all onset times, maximum over all offset times). In the Passive task, onset and offset timestamps were calculated differently by using the position raw data of the moved digit. Particularly, the range of 1-99% of the position’s gradient was chosen to mark the transient phase.

#### Individuation index in the single-finger task

Being able to voluntarily move one finger in isolation, without concomitant motion of the other fingers, is a basic pre-requisite for dexterous performance of various daily manual tasks. To quantify this ability, as expressed in flexion vs. extension, and in different digits, we computed an *Individuation Index* (II) describing the degree to which all uninstructed digits remain still during movement of a particular instructed digit. The individuation index reflects the magnitude of enslaving of the uninstructed digits at peak force of the instructed digit. Enslaving is calculated using the following equation:

Where the index j denotes the j^th^ uninstructed digit and t represents a time sample. Therefore, Forcej(t) is the force level at time sample t of the j^th^ uninstructed digit. The difference between this force and the corrected baseline force 𝑓_0_set at 0 N was summed over all uninstructed digits and averaged across all times from force initiation (“onset”, t=0) to termination (“offset”, t=T). In general, peak forces and enslaving values present a positive linear relationship, with increased instructed-digit force accompanied by increased forces generated in uninstructed digits. The Individuation index of each digit and direction was calculated as the negative log of the slope of the corresponding linear regression line (**Fig. 1E**). Hand-level indexes were obtained by averaging the digits’ indexes.

#### Normalized finger strength

To be able to compare strength impairment of different subjects, which had varying pre-morbid strength, we computed a normalized measure of non-individuated finger strength in the following manner: raw maximal finger forces (MVFs) were averaged across two assessments (see Experiment protocols section) and divided by the MVF of the unaffected hand, which was obtained in the first assessment. Due to technical issues the raw MVF data of the unaffected hand of one patient (patient #0102) were unavailable and thus were imputed using the average of unaffected hand forces of patients with the same side of limb paresis. Visual inspection confirmed that this imputation did not produce outlier data. Hand-level normalized fingers strength measure was obtained by averaging across digits.

#### Impairment in stroke patients

Comparison between stroke patients and age-matched controls revealed lower individuation capacity, lower strength, and abnormal direction-specific synergy patterns in the former group. To quantify this disadvantage, a hand-level “*Impairment*” measure (in percent) was computed for individuation index and for strength, reflecting normalized differences between patients’ values and controls’ median value. To address a potential bias of handedness, the impairment was first calculated separately for the left and right hands and then combined into a complete measure:

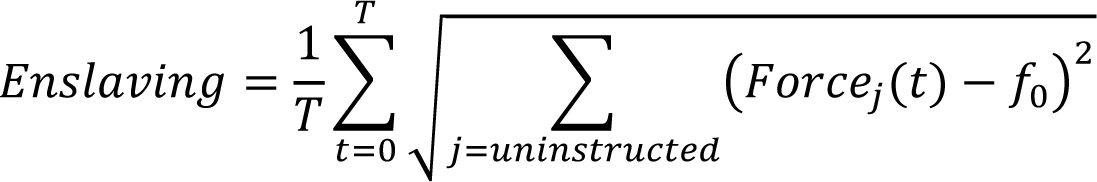

Where X may represent either individuation index or strength.

#### Accuracy and deviations in the multi-finger task

Accuracy in the multi-finger task is another key element in digit dexterity which was tested in this study. It was quantified by examining the multi-dimensional Euclidian distance between the actual force and the projection of the produced force onto the target trajectory (**Fig. 1G**). This total measurement, taking into account all digits, is called “DeviationT” and is calculated using the following equation:

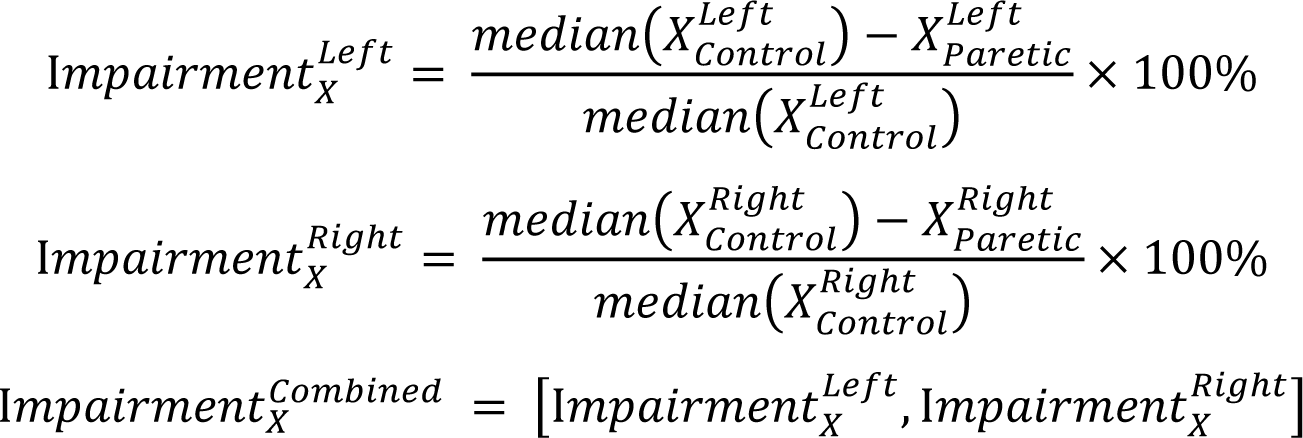

In addition, “DeviationT” can be divided into its two components: (1) “DeviationI” which quantifies the ability to generate the desired force in the instructed digit(s) and (2) “DeviationU” which quantifies the ability to avoid generating forces in the uninstructed digit(s).

“DeviationI” was calculated as follows:

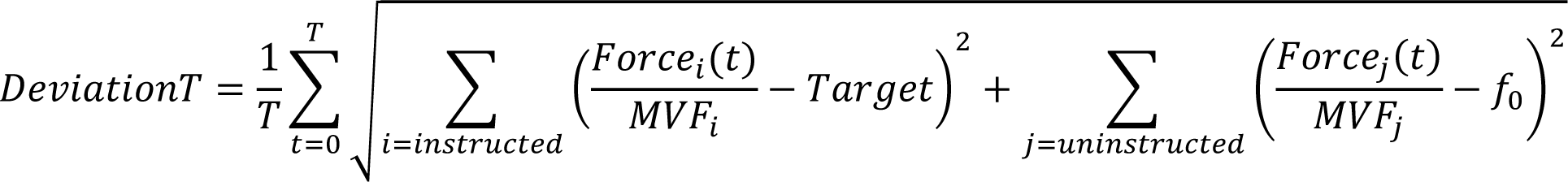

Where the index i denotes the i^th^ instructed digit and t represents a time sample. Therefore, Forcei(t) is the force level at time sample t of the i^th^ instructed digit. This force was normalized by the corresponding MVF value. The difference between this normalized force and the target set at 25% was summed over all instructed digits and averaged across all times from force initiation (“onset”, t=0) to termination (“offset”, t=T).

“DeviationU” was calculated as follows:

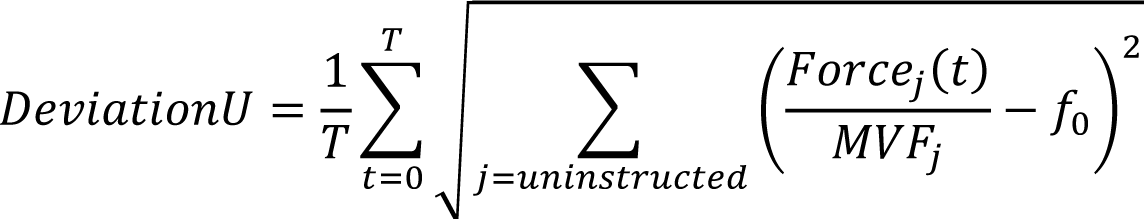

Where the index j denotes the j^th^ uninstructed digit and t represents a time sample. Therefore, Forcej(t) is the force level at time sample t of the j^th^ uninstructed digit. This force was normalized by the corresponding MVF value. The difference between this normalized force and 𝑓_0_ set at 0 was summed overall uninstructed digits and averaged across all times from force initiation (“onset”, t=0) to termination (“offset”, t=T).

#### Enslaving in the passive task

Coupling due to anatomical properties of soft tissues in the hand impose constraints on the ability to exert individuated finger movements. To isolate the effect of mechanical factors from the impact of neural control, we assessed the isometric forces generated by the unmoved digits during passive movements of each single digit by the robot. The measure of “*Enslaving*”, introduced earlier in the explanation of the single-finger individuation task, is suitable also to quantify the involuntary involvement of the unmoved digits. “*Enslaving*” in the passive task was calculated using the following equation:

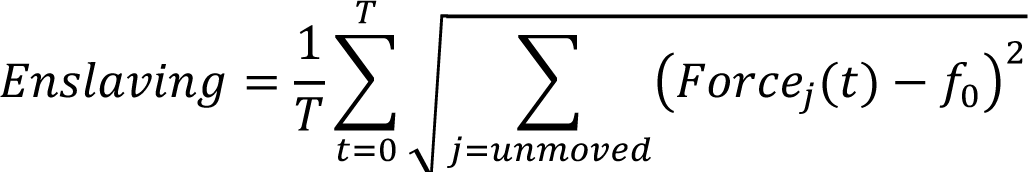

Where the index j denotes the j^th^ unmoved digit and t represents a time sample. Therefore, Forcej(t) is the force level at time sample t of the j^th^ unmoved digit. The difference between this force and the corrected baseline force 𝑓_0_ set at 0 N was summed over all unmoved digits and averaged across all times from force initiation (“onset”, t=0) to termination (“offset”, t=T).

#### Brain imaging and voxel-based lesion-symptom mapping

Follow-up CT/T1-MRI brain scans (24 CT scans, 8 T1-MRI scans) (median, IQR: 30 [7, 45] days from stroke onset) of the stroke patients were carefully examined by a physician experienced in the analysis of neuroimaging data (author SOG) to ensure that lesion boundaries were clear and traceable and that the scans present stable pattern of tissue damage without a mass effect from residual edema. Lesions were manually delineated using MRIcron software^70^ and then were registered to MNI space (MRI single subject template) at voxel-resolution of 2 x 2 x 2 mm^71^ using enantiomorphic normalization^72^ and symmetric diffeomorphic registration algorithms^73^ implemented in ANTsR package^74^ and customized R scripts. Due relatively small sample size of the cohort, all normalized lesion maps obtained from which involved the right hemisphere (n = 14) were flipped to the left hemisphere^75^. Voxel-based lesion-symptom mapping (VLSM)^13^ was then applied to the normalized binary lesion maps to identify voxels where damage had a significant impact on (1) finger individuation capacity, as expressed by the individuation index (II) in flexion and in extension directions, and (2) finger strength, as expressed by the normalized finger strength measure. Specifically, we applied Brunner-Munzel test^76^ calculated using 20,000 permutations of the data due to small sample size^77^, as implemented in LESYMAP R package^78^. Only voxels lesioned in 15% or more of the patients were included in the analysis. Family-wise error rate (FWER) was controlled using the false discovery rate (FDR) method^79^ with p-value < 0.05. Due to the relatively small sample size, we also reported voxels that did not survive these corrections, but instead survived a more lenient criterion of p < 0.01, similarly to previous reports^14–18^. To explore the extent of anatomical overlap between brain voxels implicated in finger-individuation and strength in each direction (flexion, extension), we employed VLSM conjunction analysis contrasting brain voxels that were selectively associated with either flexion or extension, and brain voxels without such direction selectivity (i.e., voxels in which the existence of damage affected performance in both flexion and extension). To unravel brain structures implicated in each measure and movement direction, significant voxels emerging from the VLSM analysis were overlayed on a set of atlas-based pre-defined regions of interest (ROIs): grey-matter ROIs based on the *Brainnetome* atlas^80^; corticospinal tracts based on the *Sensorimotor Area Tract Template* (SMATT)^64^; the cortico-reticular pathway (CRP) based on a recently published template^65^; trajectories of association tracts based on the *XTRACT* atlas and template^81^. Only voxels belonging to clusters of more than 5 adjacent significant voxels are reported.

Since significant voxel clusters were overlayed on brain structures derived from different atlases, the reported percentages may exceed 100%, due to inter-atlas overlap.

## Acknowledgments

This was supported the Israel Science Foundation (ISF) Grant 1634/19 (FM) and United States - Israel Binational Science Foundation (BSF) Grant 2021323 (FM).

## Figure Legend

## Supplementary information

### Experiment 1 – individuation index and strength at the individual digit level

The individuation index mean (±SE) values [a.u., the slope of the regression line is unitless] for the 5 digits and the whole hand in the flexion direction were: 2.82 (±0.11) for the thumb, 2.93 (±0.14) for the index, 2.58 (±0.08) for the middle, 1.50 (±0.15) for the ring, 1.36 (±0.12) for the little and 2.24 (±0.09) for the whole hand. In comparison, the corresponding values in the extension direction were: 2.30 (±0.16) for the thumb, 1.68 (±0.09) for the index, 1.45 (±0.09) for the middle, 0.88 (±0.13) for the ring, 0.92 (±0.15) for the little and 1.45 (±0.07) for the whole hand.

The strength mean (±SE) values [N] for the 5 digits and the whole hand in the flexion direction were: 18.02 (±0.47) for the thumb, 18.83 (±0.21) for the index, 18.39 (±0.39) for the middle, 16.93 (±0.6571) for the ring, 16.17 (±0.88) for the little and 17.67 (±0.45) for the whole hand. In comparison, the corresponding values in the extension direction were: 7.51 (±0.42) for the thumb, 9.26 (±0.49) for the index, 9.39 (±0.37) for the middle, 7.85 (±0.53) for the ring, 6.12 (±0.49) for the little and 8.03 (±0.32) for the whole hand. Strength was significantly lower for extension compared to flexion direction.

### Experiment 1 – performance of the multi-finger task

During the multi-finger individuation task, participants performed 25 possible finger combinations (“chords”; divided into 3 types: 10 two-finger, 10 three-finger and 5 four-finger), in which they instructed to reach toward a force level of 25% of MVF with the instructed fingers, in each direction, while keeping the other uninstructed fingers as immobile as possible. The ability to generate accurate multi-finger force combinations is separated into two aspects – force control, calculated as the deviation of the instructed fingers from the target (DeviationI, see Methods), and enslaving, calculated as the deviation of the uninstructed fingers from zero (DeviationU). These deviations were combined into a complete measurement taking into consideration all fingers (DeviationT). The lower these deviations are, the better the accuracy capability.

The DeviationI mean (±SE) values [a.u.] for the 3 chord types in the flexion direction were: 0.07 (±0.002) for the two-finger chords, 0.13 (±0.004) for the three-finger chords and 0.17 (±0.005) for the four-finger chords. In comparison, the corresponding values in the extension direction were: 0.10 (±0.005) for the two-finger chords, 0.15 (±0.006) for the three-finger chords and 0.19 (±0.009) for the four-finger chords.

The DeviationU mean (±SE) values [a.u.] for the 3 chord types in the flexion direction were: 0.08 (±0.0050) for the two-finger chords, 0.08 (±0.005) for the three-finger chords and 0.05 (±0.005) for the four-finger chords. In comparison, the corresponding values in the extension direction were: 0.1004 (±0.0081) for the two-finger chords, 0.11 (±0.011) for the three-finger chords and 0.08 (±0.010) for the four-finger chords.

Altogether, the DeviationT mean (±SE) values [a.u.] for the 3 chord types in the flexion direction were: 0.11 (±0.005) for the two-finger chords, 0.16 (±0.006) for the three-finger chords and 0.18 (±0.005) for the four-finger chords. In comparison, the corresponding values in the extension direction were: 0.15 (±0.006) for the two-finger chords, 0.21 (±0.006) for the three-finger chords and 0.22 (±0.008) for the four-finger chords.

### Experiment 2 – enslaving due to passive coupling at the individual digit level

In the *ROM* block, we examined the passive coupling on the digit-level (averaging enslaving across ROM ranges), the ROM-level (averaging enslaving across digits) and the hand-level (averaging enslaving across both ROM ranges and digits). On the digit-level, the enslaving mean (±SE) values [N] for the 5 digits in the flexion direction were: 0.20 (±0.02) for the thumb, 0.27 (±0.03) for the index, 0.30 (±0.04) for the middle, 0.35 (±0.04) for the ring and 0.28 (±0.02) for the little. In comparison, the corresponding values in the extension direction were: 0.0977 (±0.01) for the thumb, 0.19 (±0.02) for the index, 0.19 (±0.01) for the middle, 0.27 (±0.02) for the ring and 0.17 (±0.01) for the little.

On the ROM-level, the enslaving mean (±SE) values [N] for the 4 ROM ranges in the flexion direction were: 0.24 (±0.02) for 20% ROM, 0.27 (±0.03) for 40%, 0.29 (±0.03) for 60% and 0.32 (±0.03) for 80%. In comparison, the corresponding values in the extension direction were: 0.18 (±0.01) for 20% ROM, 0.17 (±0.01) for 40%, 0.18 (±0.01) for 60% and 0.21 (±0.01) for 80%.

In the *Velocity* block, we examined the passive coupling on the digit-level (averaging enslaving across velocities), the velocity-level (averaging enslaving across digits) and the hand-level (averaging enslaving across both velocities and digits). On the digit-level, the enslaving mean (±SE) values [N] for the 5 digits in the flexion direction were: 0.19 (±0.01) for the thumb, 0.26 (±0.02) for the index, 0.29 (±0.03) for the middle, 0.37 (±0.03) for the ring and 0.30 (±0.02) for the little. In comparison, the corresponding values in the extension direction were: 0.12 (±0.01) for the thumb, 0.19 (±0.02) for the index, 0.23 (±0.02) for the middle, 0.30 (±0.02) for the ring and 0.16 (±0.02) for the little.

On the velocity-level, the enslaving mean (±SE) values [N] for the 4 velocities in the flexion direction were: 0.29 (±0.02) for 0.05 m/s, 0.27 (±0.02) for 0.1 m/s, 0.28 (±0.02) for 0.15 m/s and 0.31 (±0.03) for 0.2 m/s. In comparison, the corresponding values in the extension direction were: 0.19 (±0.01) for 0.05 m/s, 0.19 (±0.01) for 0.1 m/s, 0.19 (±0.01) for 0.15 m/s and 0.2 (±0.01) for 0.2 m/s. On the hand-level, the enslaving mean (±SE) value [N] for the whole hand in the flexion direction was 0.29 (±0.02). In comparison, the corresponding value in the extension direction was 0.19 (±0.01). Overall, enslaving was significantly lower for extension compared to flexion direction.

Furthermore, we were able to replicate our results (i.e., finger individuation and strength) from Experiment 1. Here, the individuation index mean (±SE) values [a.u.] for the 5 digits and the whole hand in the flexion direction were: 2.65 (±0.09) for the thumb, 2.59 (±0.08) for the index, 2.4 (±0.12) for the middle, 1.59 (±0.13) for the ring, 1.43 (±0.12) for the little and 2.14 (±0.07) for the whole hand. In comparison, the corresponding values in the extension direction were: 2.15 (±0.11) for the thumb, 1.54 (±0.16) for the index, 1.27 (±0.13) for the middle, 0.75 (±0.13) for the ring, 0.81 (±0.17) for the little and 1.30 (±0.10) for the whole hand. Once again, individuation was significantly lower for extension compared to flexion direction. This can be seen in two-way (direction vs digit) RM-ANOVA, which showed significant effects for the direction factor (𝐹_1,18_ = 96.22, 𝑝 = 1.2 × 10^−8^) and the digit factor (𝐹_4,72_ = 48.32, 𝑝 = 1.11 × 10^−1^^9^), as well as an interaction between the two (𝐹_4,72_ = 4.56, 𝑝 = 0.0024).

The strength mean (±SE) values [N] for the 5 digits and the whole hand in the flexion direction were: 19.05 (±0.39) for the thumb, 19.75 (±0.53) for the index, 19.74 (±0.30) for the middle, 19.04 (±0.50) for the ring, 17.5914 (±0.5940) for the little and 19.0290 (±0.3138) for the whole hand. In comparison, the corresponding values in the extension direction were: 7.53 (±0.37) for the thumb, 7.62 (±0.3653) for the index, 8.27 (±0.36) for the middle, 6.70 (±0.40) for the ring, 5.78 (±0.48) for the little and 7.18 (±0.29) for the whole hand. Once again, strength was significantly lower for extension compared to flexion direction. This can be seen in two-way (direction vs digit) RM-ANOVA, which showed significant effects for the direction factor (𝐹_1,19_ = 746.36, 𝑝 = 1.03 × 10^−1^^6^) and the digit factor (𝐹_4,76_ = 12.37, 𝑝 = 8.46 × 10^−8^), though no interaction between the two (𝐹_4,76_ = 0.57, 𝑝 = 0.68).

### Experiment 3 - individuation index and strength at the individual digit level

The stroke group’s individuation index mean (±SE) values [a.u.] for the 5 digits and the whole hand in the flexion direction were: 1.62 (±0.18) for the thumb, 1.34 (±0.20) for the index, 1.02 (±0.18) for the middle, 0.52 (±0.13) for the ring, 0.07 (±0.18) for the little and 0.91 (±0.15) for the whole hand. In comparison, the corresponding values in the extension direction were: 1.2387 (±0.1627) for the thumb, 0.31 (±0.19) for the index, 0.47 (±0.18) for the middle, -0.08 (±0.13) for the ring, 0.30 (±0.16) for the little and 0.44 (±0.12) for the whole hand.

The control group’s individuation index mean (±SE) values [a.u.] for the 5 digits and the whole hand in the flexion direction were: 2.90 (±0.08) for the thumb, 2.70 (±0.11) for the index, 2.25 (±0.16) for the middle, 1.32 (±0.09) for the ring, 1.34 (±0.17) for the little and 2.10 (±0.07) for the whole hand. In comparison, the corresponding values in the extension direction were: 2.27 (±0.165 for the thumb, 1.72 (±0.15) for the index, 1.57 (±0.12) for the middle, 0.70 (±0.15) for the ring, 1.07 (±0.1700) for the little and 1.48 (±0.08) for the whole hand.

The stroke group’s strength mean (±SE) values [N] for the 5 digits and the whole hand in the flexion direction were: 12.60 (±1.00) for the thumb, 12.11 (±0.97) for the index, 12.41 (±0.93) for the middle, 10.74 (±0.96) for the ring, 8.54 (±1.01) for the little and 11.28 (±0.89) for the whole hand. In comparison, the corresponding values in the extension direction were: 3.85 (±0.46) for the thumb, 3.39 (±0.45) for the index, 3.96 (±0.54) for the middle, 2.48 (±0.40) for the ring, 3.13 (±0.37) for the little and 3.36 (±0.41) for the whole hand.

The control group’s strength mean (±SE) values [N] for the 5 digits and the whole hand in the flexion direction were: 17.47 (±1.01) for the thumb, 17.60 (±1.02) for the index, 17.02 (±1.06) for the middle, 15.08 (±1.18) for the ring, 15.20 (±1.29) for the little and 16.47 (±1.05) for the whole hand. In comparison, the corresponding values in the extension direction were: 6.2017 (±0.6961) for the thumb, 6.42 (±0.60) for the index, 6.36 (±0.40) for the middle, 4.84 (±0.43) for the ring, 4.73 (±0.53) for the little and 5.71 (±0.41) for the whole hand.

